# Probing the dependence of transcription factor regulatory modes on promoter features

**DOI:** 10.1101/2024.05.30.596689

**Authors:** Sunil Guharajan, Vinuselvi Parisutham, Robert C. Brewster

**Affiliations:** Department of Systems Biology, University of Massachusetts Chan Medical School, Worcester, MA 01605; Department of Microbiology and Physiological Systems, University of Massachusetts Chan Medical School, Worcester, MA 01605

## Abstract

Transcription Factors (TFs) are often classified as activators or repressors, yet these context-dependent labels are inadequate to predict quantitative profiles that emerge across different promoters. The regulatory interplay between a TFs function and promoter features can be complex due to the lack of systematic genetic control in the natural cellular environment. To address this, we use a library of *E. coli* strains with precise control of TF copy number. We measure the quantitative regulatory input-output function of 90 TFs on synthetic promoters that isolate the contributions of TF binding sequence, location, and basal promoter strength to gene expression, uncovering TF specific regulatory principles. We infer that many of these TFs function by stabilizing RNA polymerase at the promoter, a property we see for both activating and repressing TFs. We develop a thermodynamic model that predicts stabilizing TFs have a specific quantitative relationship with promoters of differential strength. We test this prediction using synthetic promoters spanning over 100-fold range in basal expression levels and confirm that stronger promoters have lower fold-change for stabilizing TFs, whereas non-stabilizing TFs do not exhibit this relationship, indicating a conserved mechanism of transcription control across distinct TFs. This work demonstrates that understanding the intrinsic mechanisms of TF function is central to decoding the relationship between sequence and gene expression.

## Introduction

For decades, TFs have been classified as either activators or repressors based on their net regulatory function on a promoter. However, as we have learned more about the complex interactions TFs have with transcriptional machinery, it has become increasingly clear that these labels are highly context-dependent and rely on specific details of each promoter - such as which other TFs regulate it and where they bind. Any individual TF may regulate dozens of different promoters, each with a distinct genetic context and each of those that may take cues from physiological signals sensed or imposed upon the cell. As a result, these coarse labels of “activator” or “repressor” have proved to be less valuable when trying to predict how a TF will impact the expression of a gene in contexts where the action of that TF is not already known, such as under different stresses or mutations of the promoter region. For that reason, being able to characterize TFs based on their interactive properties with polymerase is a promising approach to developing a predictive level understanding of regulatory circuit design.

Bacterial transcription is a multi-step process and the net expression from a promoter relies on the rate at which RNA polymerase can progress through them [1–3]. These steps include the binding of polymerase at the promoter to form the closed complex, promoter melting to form the open complex and finally the initiation of transcription. TFs may regulate expression by altering the rate of one or more of these steps [4–14]. As such, many combinations of regulatory mechanisms can lead to the same apparent change in expression to a specific promoter. Promoter sequence determines the basal rate at which RNAP can progress through the steps of transcription [15, 16] and thus the interplay between the regulatability of a promoter by a TF and the kinetic steps of RNAP binding and promoter clearance is fundamental to understanding how control of endogenous promoters is achieved.

Many features of the local sequence and cellular state impact the degree to which a promoter is regulated. Crucial factors such as the presence of TF binding sites adjacent to the promoter, their relative position to the promoter, and their sequence are all genetically encoded determinants of regulation. Furthermore, the state of the cell will control the identity, availability and activity of factors that may bind these sites. At a basic level, each of these factors may alter two fundamental aspects of the TFs ability to regulate a promoter: the TFs propensity to be bound at a specific binding site and the TFs physical effect on the transcription process that it exerts when bound. However, the contributions of TF binding sequence, TF binding location, and the basal strength of the regulated promoter play fundamental roles in realizing the net regulatory outcome of a TF on gene expression (Fig. 1A), yet the convolution of each of these important sequence features in natural promoters makes isolating their individual contributions and mechanisms of action challenging.

**Figure 1:**
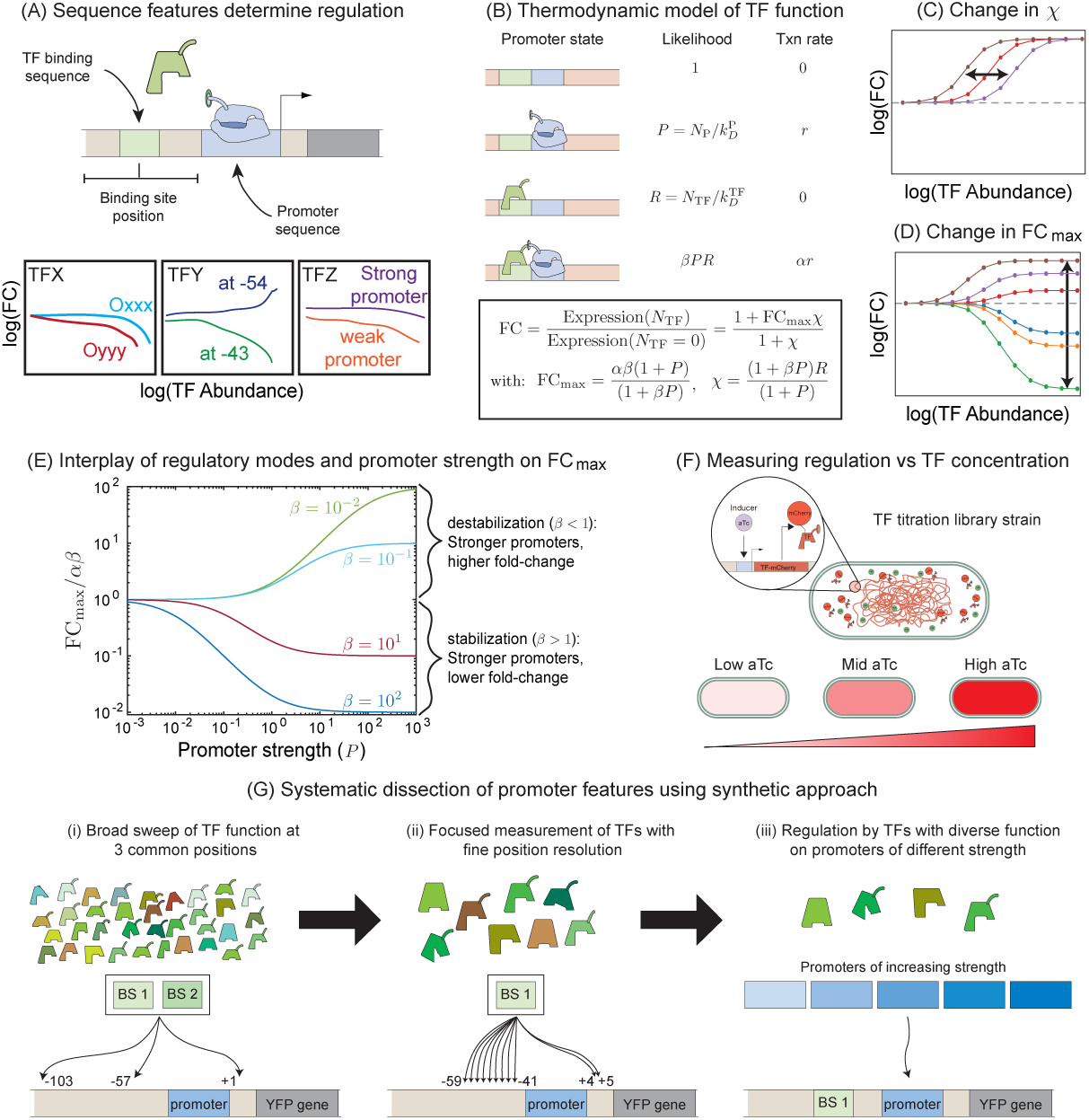
Measuring regulation of *E. coli* TFs systematically. (A) TF regulatory function depends on binding sequence, binding position on the promoter, and promoter identity. (B) Simple thermodynamic model to characterize the regulatory role of TFs (C-D) Model predictions depend on two primary features: occupancy (C) and regulatory function (D). (E) Role of promoter, *α* and *β* in setting the max regulatory effect of a TF, FC_max_. (F) Systematic TF expression library enables precise control of individual TF levels. (G) Overview of experiments here to test the role of position, binding sequence and promoter identity in setting the regulatory response functions of *E. coli* TFs.

There is a well-established relationship between binding site sequence and TF occupancy that has been elucidated in models of sequence-based binding affinity[17–20] informed by high-throughput techniques both *in vitro* [21–27] and *in vivo* [20, 28–30]. TF binding location on the promoter is a well-established determinant of regulatory function in bacteria with the magnitude of regulation typically showing periodicity following the 10.5 bp helicity of double-stranded DNA structure [31–36].However, there is also evidence from quantitative investigations of TF function that demonstrate certain sequences affect regulation beyond merely impacting TF occupancy at the site [37–41].

A simple model of TF function, outlined in Fig. 1B, explicitly separates the contributions of TF function and TF occupancy. Here the system is confined to the effects of only a single TF which regulates expression through two mechanisms: regulation of the stability of RNAP at the promoter (parameterized as *β*) and regulation of the rate of transcription from promoter-bound RNAP (parameterized as *α*). From this model, it can be seen that the regulation exerted by any TF can be encapsulated into an effective TF occupancy, *χ*, and the net regulatory effect of the TF, FC_max_, which make unambiguous predictions for the “response function” of a promoter to the TF (Fig. 1C,D). However, one interesting prediction of this model is how TFs will interact with different promoters, parameterized by promoter strength *P*. TFs regulate many distinct target genes with different promoter sequences [42], furthermore, physiological changes will regulate the overall rate of transcription in the cell by controlling polymerase availability [43–46]. As shown in Fig. 1E, this model predicts that the relationship between the regulatory effect of the TF (FC_max_) and the strength of the promoter regulated (*P*) that depends on the TFs usage of the stability mechanism, *β* in our model.

To assess the spectrum of regulatory responses from a diverse array of TFs in *E. coli*, we used a library of strains where the concentration of a given TF in the cell is titratable with the small molecule inducer, aTc [41]. This system allows for precise control and measurement of TF copy number: the TF is fused to the mCherry open reading frame through a linker sequence, which allows us to measure the input level of TF in flow cytometry at the single-cell level (see Fig. 1F) [14, 41]. The decoupling of the TFs in the library strains from their native genetic context allows us to induce expression of individual regulators in *E. coli* between several copies per cell up to hundreds or thousands (see Methods Section and ref. [41]), allowing us to separate changes in gene expression by the direct effect of the TF on the promoter. Armed with quantitative control of TF induction, our goal was to systematically dissect the role of *cis*-acting determinants of the promoter on each of the individual TFs regulatory ability - a broad overview of which is illustrated in Fig. 1G. We initially profiled a vast array of regulators by targeted selection of TF binding sequence and location (Fig. 1Gi), followed by a more detailed exploration of the position landscape of regulation for select TFs (Fig. 1Gii). Finally, we aimed to use the information on the regulatory properties of these TFs - particularly their utilization of stabilization - to predict how these TFs would function on promoters of different strengths (Fig. 1Giii).

## Results

### Assessing the Isolated Regulatory Function of 90 *E.coli* TFs

To comprehensively assess the regulatory function for a broad number of E.coli TFs, we selected a total of 90 TFs from the TF titration library [41] and used the RegulonDB database [47] to identify known binding sequences of each TF. From this list of binding sites, we selected 2 *−* 3 sequences for each TF based on several criteria: strong evidence of binding in RegulonDB and the target promoter is *σ*^70^. In total, 180 TF binding sequences were selected (full list in supplemental materials). Each binding site sequence was cloned into 3 different binding positions relative to an otherwise unregulated *σ*^70^ promoter on a low copy number plasmid (Fig. 2A). In total, 540 synthetic circuits with distinct regulatory architectures were cloned. By independently controlling both the binding position and sequence of each TF in a designed regulatory circuit we sought to investigate the role of these features on the quantitative input-output regulatory function of a collection of diverse TFs. The data for Fold-Change as a function of TF copy number is shown in Fig. 2B. Each curve is colored by the maximum level of regulation achieved, from repressive (blue) to activating (red).

**Figure 2:**
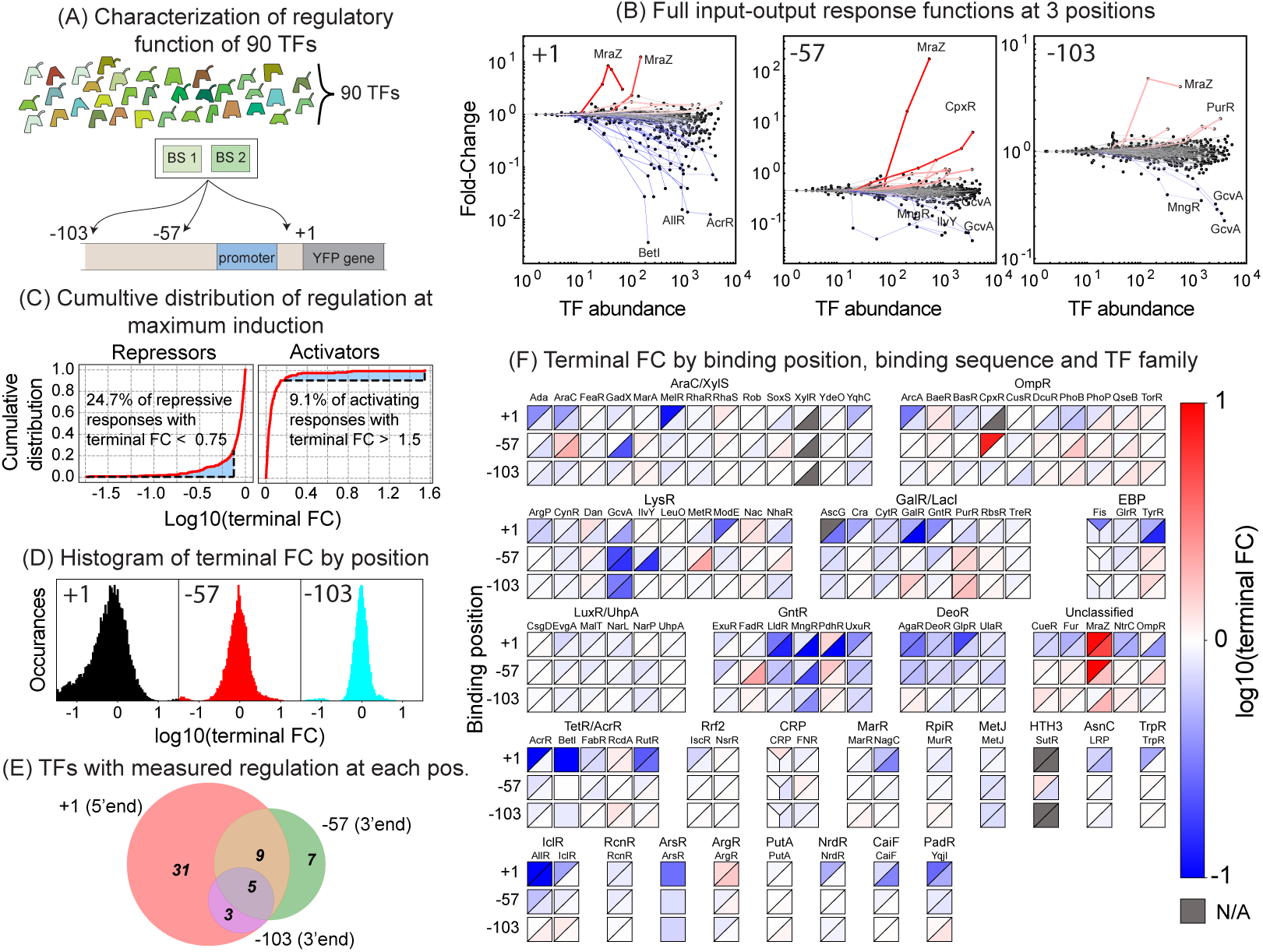
Measuring the Response Function of 90 TFs. (A) 90 *E.coli* TFs encompassing regulation of several key cellular functions were profiled on synthetic circuits to isolate the fundamental response function. Each regulator was assayed at 3 binding locations: immediately downstream (+1) relative to the promoter, a proximal promoter position (*−*57) associated with activation, and a distal position (*−*103). An average of 2 binding sites per TF were selected as described in the Methods Section and cloned into each of the three positions in the circuits. (B) Measured response functions across 531 regulatory circuits. Note the presence of most repression at the +1 binding location, consistent with the intuition of regulation immediately downstream. Activating interactions are more common at position *−*57. (C) Cumulative Distribution Function (CDF) for the response function. A response function was classified as activating if the terminal FC value was above a threshold 1.5-fold and repressing if the FC was below 0.75. Bounded, shaded regions represent the area (fraction of response functions) classified into activation and repression. (D) Density function for the maximal FCs for each of the three binding positions. A kernel density estimate was used to estimate the distribution on the interval (*−∞*,+*∞*) with values *>*= *|*1.5*|* truncated from the plot. (E) Position overlap of regulators with the distal position (*−*103) demonstrating activity for only 8 tested TFs.(F) Quantitative Atlas depicting the terminal FC values for each of the TFs at their respective biding locations and sequences. The color scheme denotes blue for repression and red for activation. Note the area of a square for a given TF is partitioned based on the number of sequences measured for the regulator.

From the measured regulatory response functions for the 540 synthetic circuits, we removed 9 response functions with spurious fold-change profiles, considering only 531 synthetic circuits for subsequent analysis where we find a spectrum of regulatory effects. We determined a cutoff for classifying a specific interaction as activating, repressing, or no clear regulatory effect (see Fig. 2C). This was done by comparing the measured “terminal fold-change” (calculated as the average fold-change across the two highest induction conditions) to the measured fold-change in expression of the same gene when the binding site was removed (see Fig. S1). We find that across all binding locations, binding sequences and TF identities, approximately 20% of our measured response functions (104 out of 531) had clear regulatory signals outside the bounds of the no-binding-site control. As expected, the proximal binding site +1 confers the largest number of regulatory interactions with 69 (65 repressing, 4 activating), while *−*57 has 26 instances (18 repressing, 8 activating) and *−*103 has just 10 (4 repressing, 6 activating). The accompanying histograms of terminal fold-change separated by binding position are shown in Fig. 2D. A Venn diagram showing the overlap of TFs that are found to function at the three positions is shown in fig. 2E. In total, we find 55 of the 90 TFs has at least one measured regulatory interaction. The remaining 35 TFs that did not pass our threshold could be due to several reasons. We believe the primary cause to be the relatively few binding positions evaluated in these measurements combined with a strict threshold for calling regulatory interactions. For example, SoxS, a TF whose regulatory profile we examine in the next section and shows strong regulation at many positions, fails to pass the threshold at these three binding positions. As such, our filter for distinguishing regulatory interactions is comparatively strict and favors false negatives over false positives. Furthermore, we have seen strict position sensitivity for a handful of other TFs in the past, for example, we recently found the TF ZntR has no regulatory activity at +1 but strong repressive behavior at +5 [41]. However, other factors likely play a role for some TFs; the chosen binding site sequences may be incorrect or low affinity, the TFs may not be active in our growth condition, the fusion with mCherry may oblate some TF’s regulatory function, or some TFs may have an intrinsically weak regulatory effect.

In Fig. 2F, we show an overview of the terminal fold-change of each TF for all positions and sequences tested. The data is grouped according to TF families: the rows represent the 3 binding positions ordered from proximal to distal and the color map represents the measured terminal Fold-Change for each TF (the average Fold-Change at the 2 highest induction conditions). The area of the squares is partitioned based on the number of binding sequences assayed for that TF, as such each section of a square represents measurements of regulation for a given TF at a given position for different binding sequences. Overall, we see the most regulatory effects at the binding position +1 which are dominated by repressive interactions. In many cases, model TFs have been shown to regulate at +1 through a steric hindrance between the TF and RNA polymerase at the promoter, this has been demonstrated for TFs with diverse functions for example in canonical repressors such as LacI [48–50] and TetR [51, 52], as well as activators such as CRP and CpxR [14, 53]. While repression is very common at +1 we also we see the MraZ TF with significant activation at +1, suggesting potential interactions with RNAP that are not exclusively inhibitory at this position.

The data across positions provides a quantitative snapshot of the isolated TF function for TFs across several superfamilies. For instance, we find that the MngR TF, regulates with almost consistent Fold-Change across both sequence and positions immediate and distal to the promoter, while PdhR seems to have activating and repressing interactions depending on the sequence it binds to. Furthermore, the TFs UxuR and ExuR are known paralogs that regulate hexuronate metabolism and bind interchangeably to their cognate sequences as the TFs share a conserved DNA binding motif [54, 55]. Here, we demonstrate that while both TFs directly repress transcription, the quantitative activity of UxuR is noticeably stronger than its counterpart at the downstream position, suggesting either an increased binding affinity to the sequence or a stronger maximal regulatory effect. Taken together, assessing the isolated response function of 90 *E.coli* TFs uncouples position and sequence activity profiles for several factors, bridging the gap between qualitative promoter binding information and functional regulatory output for the first time for several of these factors.

### Delineating the role of TF binding sequence and location on kinetic regulatory properties of the TF

Analysis of the 531 response curves reveals a rich regulatory compendium for a subset of TFs that function across the 3 different locations tested as well as multiple binding sequences. This provided us with a means to determine how quantitative control over transcription at distal (-103), proximal upstream (-57), and proximal downstream (+1) binding and across distinct sequences is achieved by these TFs. Probing TF dependence of regulatory properties (propensity to (de)stabilize and (de)accelerate) over these tested positions and sequences would inform of us of TF-specific regulatory control – providing a rationale for more extensive characterization along either of these cis-regulatory axes. To do so, we contextualized the thermodynamic model in a Bayesian inference scheme (See Methods section for details) to extract the degree to which the profiled TF binding sequences and positions control the regulatory function (FC_max_) of 28 TFs (see Fig. 3A).

This procedure is outlined in Fig. 3B using data from positional regulation of the TFs MngR and AraC. For MngR (top two panels), the model allowing FC_max_ to change with binding position (left, green box) does not describe the data significantly better than the model where FC_max_ has a single value for all three positions (right, blue box). As such, the Bayes factor is less than 1, and the “independent model” is favored. On the other hand, AraC clearly demonstrates a case where a single value of FC_max_ will fail to describe all three curves and the corresponding Bayes factor is greater than 1, favoring the model where FC_max_ depends on the tested positions. In other words, for MngR the regulation at the binding positions measured here can be described without the need to invoke changes to the regulatory properties of the TF with position, whereas for AraC the function (its ability to (de)stabilize or (de)accelerate RNAP) must change with position.

**Figure 3:**
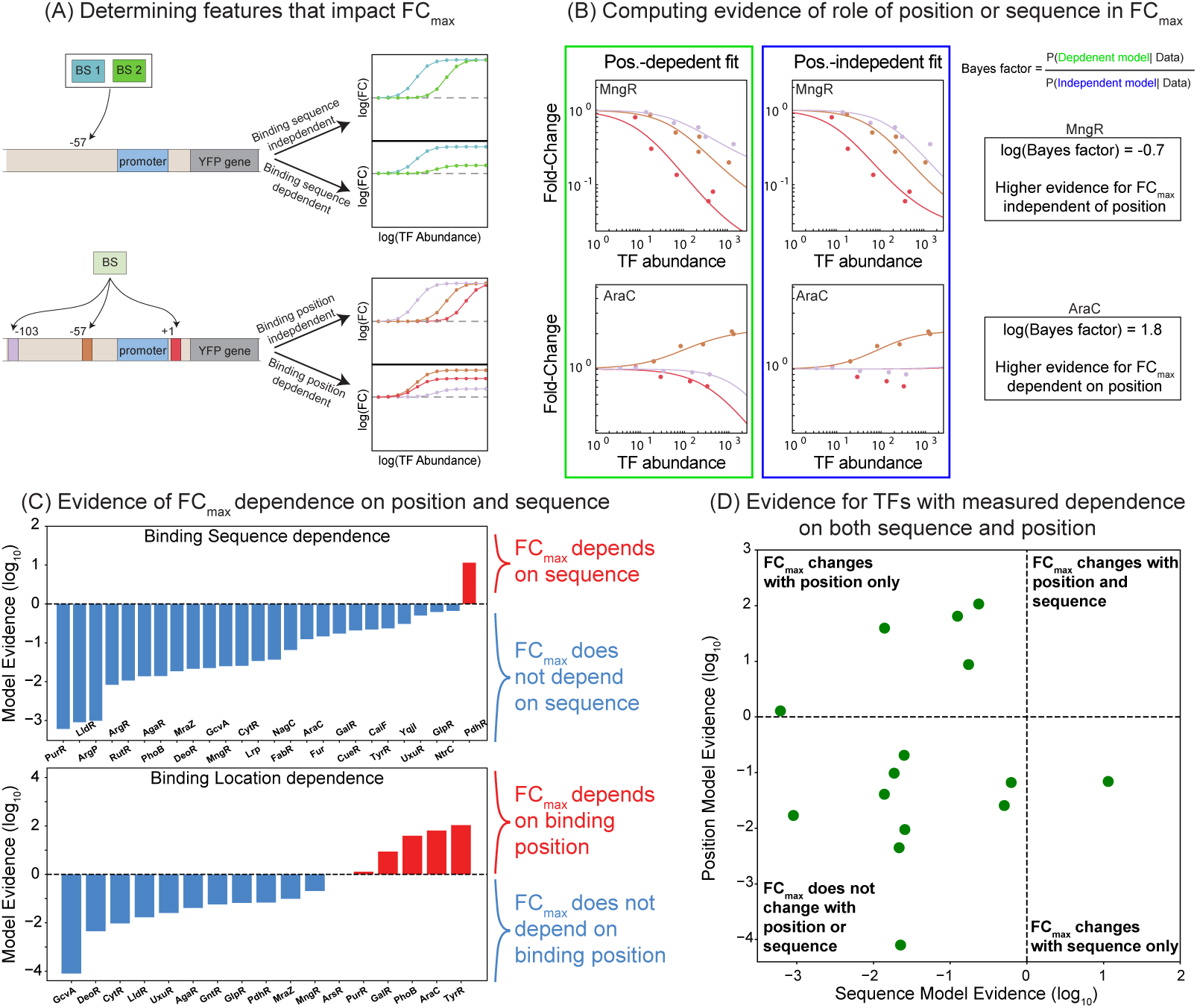
The role of TF binding sequence and position in setting TF regulatory function. (A) Gene regulatory circuits that systematically vary TF binding sequence at a fixed position, and vary TF binding position with fixed sequence allow us to assess the relative contributions of each feature to TF regulatory function (FC_max_ in the thermodynamic model). (B) Assessing evidence for the role of position in setting TF regulatory function (*α* and *β*) for TFs AraC and MngR. Left column is the thermodynamic model where FC_max_ is allowed to vary between different position response functions and right is the model with the parameter fixed. Note that MngR regulation proximal (+1) and distal (*−*103) is effectively capture by a single FC_max_ while AraC is not. (C) Model Evidence on the role of TF binding sequence for 26 TFs. As demonstrated, all the TFs with the exception of PdhR demonstrate more evidence in favor of the model where sequence is constrained (FC_max_ is fixed across the response functions). Evaluating the role of binding sequence reveals 5 TFs (PurR, GalR, PhoB, AraC, TyrR) requiring binding position to explain the data. (D) Scatter plot for 15 TFs with regulatory responses that meet the criteria for dual evaluation of position and sequence. A majority of the TFs (9 out of 15) have constant TF regulatory function (ability to stabilize and accelerate RNAP) across the sequence and positions tested.

In Fig. 3C, we calculate the Bayes factor for both position and sequence dependence of FC_max_ for each TF where we measure regulation at more than one position or sequence. The top histogram demonstrates the Bayes factor for regulation at different binding sequences for 24 TFs. We find that for each TF except one (PdhR), the data is well explained by the model where TF function (FC_max_) is independent of the tested sequences. This is consistent with our understanding of the role for sequence in contributing to TF occupancy at the promoter and suggests that sequence is decoupled from the other promoter features that materially set TF regulatory function. In contrast, for 16 TFs where multiple binding positions demonstrate regulation, we find 5 of these data sets are better described by a model where TF function depends on the tested positions (red histograms) than the independent model (blue histograms). There are 15 TFs where we measure the Bayes factor for across both feature sets and these are shown in Fig. 3D. An important caveat to our method is that in cases where the TF regulates at one position on the promoter, but shows no regulation when the binding site is moved to a different position on the promoter. This can be equally well described in the model as ablation of function (which is typically assumed) or as ablation of binding. As such, data such as this does not influence the Bayes factor. However, even without this assumption, we find that binding sequence did not typically impact function whereas binding position did for a greater subset of the TFs.

### Positional regulation of 24 TFs unveils the quantitative landscape of promoter-proximal regulation

As our statistical inference procedure confirmed that binding position was a significant determinant of TF function for certain factors, we selected 24 of the 90 TFs that showed significant regulatory activity for at least one of the three binding locations (+1, *−*57 or *−*103) or were previously attested in the literature to have position-specific regulation. We then measured their regulatory activity across 11 different binding site locations spanning the promoter for a single binding sequence. These 24 TFs belong to a diverse set of transcriptional regulators in *E. coli* that encompass multiple physiological processes in the cell including multi-drug eflux pathways (AcrR, BaeR) [56–59], regulation of the heavy-metal exposure response (CueR, CusR) [60–62], nitrogen stress (nac, NtrC) [63, 64], and various metabolic pathways (AraC, MetR, GcvA, AgaR, YqhC) [65–69] to name a few. As shown in Fig. 4A, the selected binding positions included almost every upstream region between *−*41 and *−*59 in steps of 2 base pairs (excluding *−*45, which we found to have an unintended mutation in the final cloning vector) as well as downstream positions: +4 and +5.

**Figure 4:**
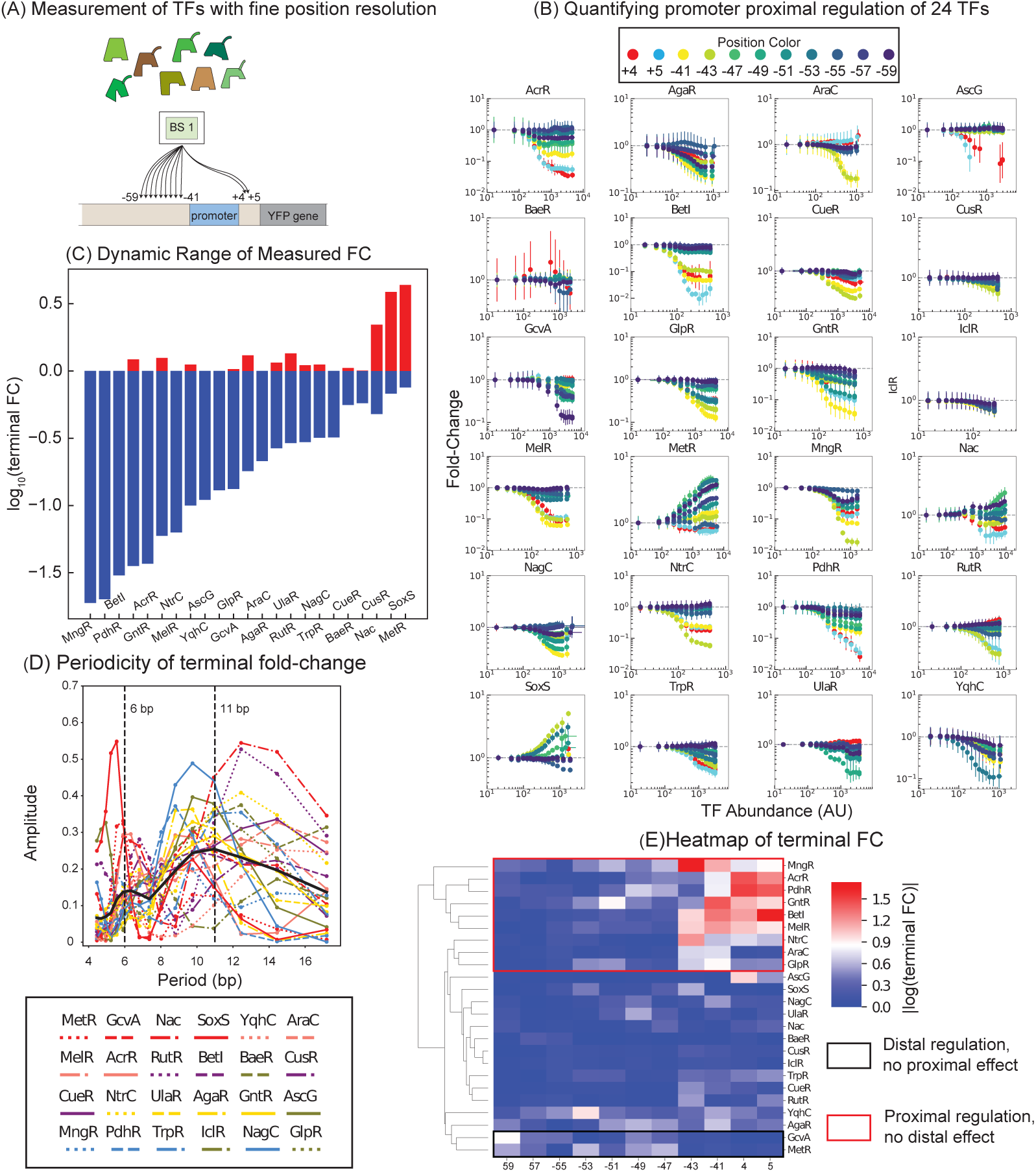
Characterizing the regulatory diversity of *E.coli* TFs across position reveals rules for proximal promoter regulation. (A) Schematic detailing the synthetic circuits used to measure 24 TFs at 11 binding positions (ranging from *−*59 bps from the TSS to +5). (B) Binding position dependence of these factors with diverse *in vivo* physiological functions. Colors represent the binding site position relative to the synthetic promoter, with 9 upstream and 2 downstream positions (11 total) measured for each TF. (C) The dynamic range of terminal Fold-Change measurements for each TF determined from position specific regulatory curves. With the exception of Nac, where parity is observed for activation and repression, all other TFs preferentially regulate as activators or repressors. (D) Periodogram of the regulatory response for all 24 TF position profiles determined using the Lomb-Scargle algorithm. The average profile shows a peak around 11bps and a smaller peak at 6bps, in line with the periodicity of DNA.(E) Unsupervised clustering of TF response function across position reveals groups of TFs that regulate with defined spatial profiles.

The full measured input-output functions for each TF are shown in Fig. 4B. As seen in the data, repression is more common than activation. This is despite several TFs acting as activators in some endogenous contexts (see Fig. S6). Across the dataset, we find one hundred-fold dynamic range when measuring repressors, with TFs such as MngR, GntR, and BetI featuring the strongest effects (Fig. 4C, Fig. S5). For the three TFs that strongly activate gene expression: SoxS, MetR, and Nac we find that the magnitude of the response does not increase by more than 6*−*fold, however, these TFs have a slightly expanded dynamic range due to their ability to repress as well as activate. For example, Nac shows a mixture of effects depending on binding position - with two-fold or more activation at positions *−*49 and *−*59 and repression of expression at downstream binding to the promoter at positions +4 and +5. We further examined the spatial regulatory patterns for periodicity; in previous studies, many TFs have been shown to obey a 10.5bp periodicity in regulatory magnitude of effect due to the length-scale set by the helicity of DNA [35, 70–72]. In Fig. 4D we calculated the spatial periodicity of the measured terminal fold-change for each TF as well as the average across all TFs (black line). We find that an 11 bp period is the dominant mode with a smaller peak at roughly half of this period (6 bp), although each individual TF has a peak *±*1 base pair from this overall average. The limited range of our measurements (19 bp) likely contributes to the noise in this decomposition. An example of the periodicity can be seen for SoxS, where activation occurs prominently at positions *−*43 and *−*53 (Fig. S5).

In Fig. 4E, we cluster the heat map of the terminal Fold-change values based on their spatial pattern of regulation. Three primary behaviors can be seen here: TFs without strong regulation, strong regulation proximal to the promoter (+5 through *−*43) and TFs with strong distal regulation (*−*49 through *−*59). As such, we do not see a simple distance-dependent rule for the strength of regulation across these measured TFs. However, a majority of the TFs that repress, do so strongly at downstream positions. This is intuitive since the downstream DNA of the promoter is involved in the transcription process, but the mechanism of this regulation is not clear; TFs may occlude polymerase binding (*β <* 1), make initiation slower (*α <* 1) or both. Taken together, these quantitative measurements at fine position resolution reveal the promoter-proximal regulatory function for diverse TFs - with differences in position function hinting at possible distinctions in the TFs ability to regulate recruitment of RNAP and initiation of transcription.

### *E.coli* TFs that repress and activate gene expression present evidence for stabilization in the regulatory response

Armed with the quantitative input-output function for a suite of regulatory factors in *E. coli*, we sought to deconvolve net regulation as a function of the TFs regulatory properties (*α* and *β*) as delineated in Fig. 1B. As TFs can alter distinct kinetic steps of the transcription process, different quantitative profiles could emerge despite similar regulatory activity at a given step of the process. The approach we use to determine the regulatory mechanisms is outlined in Fig. 5A; we re-plot the positional regulatory data as the foldchange when the TF binding site is at one position against the fold-change when that same binding site is at a different position on the promoter. Each data point on this plot represents a measurement of each of the two promoters under the same TF induction condition. We expect points in this space, which are pairs of fold-change values of the two promoters, to go from (1, 1) in the absence of the TF to 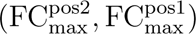 at saturating concentration of TF. The schematic in Fig. 5A shows how we expect this data to look when comparing two positions that repress (top left panel), two positions that activate (bottom right panel) and one position that activates and one that represses (top right and bottom left panels). The shape of this curve depends on a parameter *κ* = (1 + *β*_pos1_*P*)*/*(1 + *β*_pos2_*P*) which helps us assess if a given regulatory interaction is using stabilization (*β >* 1). However, the relationship is complex: if *β* is small at each position (such that *βP ≪* 1) or if *β*_pos1_ = *β*_pos2_, we expect the data to follow a straight line (black lines in Fig. 4B). On the other hand, if the data curves towards either axis, it means that *β* at the position plotted on that axis is higher than at the other position; for instance, for each blue dashed curve in Fig. 4B *β_x_ > β_y_*, and for red curves *β_y_ > β_x_*. In the case that one of the *β* values is small, the value of *β* for the other position can be inferred, as long as *βP >* 1. We have previously used this technique to infer the regulatory mechanisms of CpxR [14]. Overall, we can use this approach to find candidate TF interactions that operate through a mechanisms of stabilization with RNAP.

**Figure 5:**
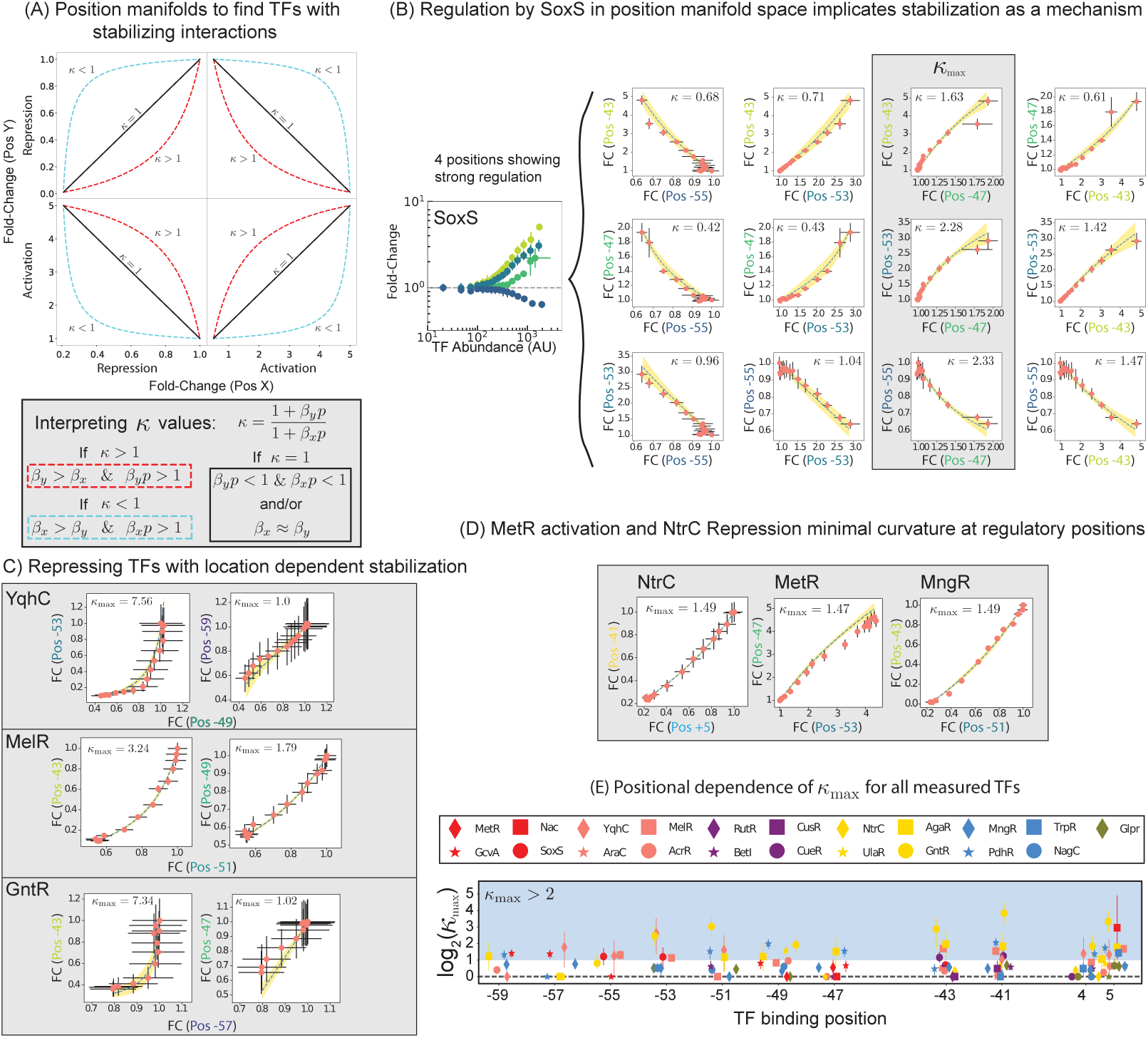
Concentration Manifold Approach Delineates Positions of Stabilizing Regulation for Select TFs. (A) Diagram of all possible outcomes of pairwise-regulatory comparisons using the manifold approach. The red curve indicates significant curvature in the data for the regulatory position in the Y-axis, and the blue curve demonstrates profiles where curvature is prominent for the position on the X-axis. Regulatory interpretation for curvature in the response function could be due to the presence of stronger stabilization relative to the reference position. (B) Pairwise Manifold Analysis for the SoxS TF across 4 positions with appreciable regulation. Positions *−*43, *−*53, and *−*55, with responses that range from mild repression to activation, have strong curvature in the regulatory response as compared to the -47 position. Cyan lines represent the manifold model curves derived from thermodynamic assumptions with shaded regions representing one standard deviation of the expected model. (C) Repression by stabilization of RNAP could potentially be in action for TFs YqhC, MelR, and GntR. (D) TFs NtrC, MetR and MngR have flat profiles across the regulatory positions, suggesting a potential absence of stabilization or stabilization at several regulatory positions. (E) Pairwise manifold analysis for 21 TFs with the fitted Kappa values at a given binding position scaled by the smallest Kappa value in the dataset. Dots represent the median kappa values and lines encompass 86*^th^* percentile of the values derived from bootstrapping. Note the predominance of the repressor GntR at both upstream and downstream regulatory positions as well as several TFs that solely repress expression. Errorbars on all manifold plots represent measurement uncertainty in the position specific fold-change

An example of the manifold analysis is shown in Fig. 5B for the SoxS regulatory response across promoters. As SoxS has previously been described in the literature to activate expression by modulating the recruitment of RNAP to the promoter, we tested the degree of curvature across the regulatory positions that showed an appreciable regulatory response (*−*43, *−*47, *−*53 and *−*55). In Fig. 5B we plot each permutation of Fold-Change at position *x* vs *y* for the 4 regulatory positions of SoxS to identify positions with significant curvature in the regulatory response. Incorporating all possible position combinations allows for a robust estimation of the curvature present in the response at a given position (see Methods Section). Within each plot we also show the measured value of *κ* for that pair of positions. When *−*47 serves as the reference position, *κ* at all other positions is maximized, making it a useful position for comparison. This is because at this position 1 + *κβ* is closest to 1 and thus we can estimate a lower bound of *β* at the other positions. We refer to these values of *κ* as *κ*_max_; these plots are highlighted in gray in Fig. 5B. The positions *−*43, *−*53 and *−*55, all have detectable curvature in the response, implying these positions all have stabilizing regulatory interactions. This is in agreement with previous studies on the regulatory mechanisms of SoxS which are known to recruit RNAP to the promoter [73, 74]. Interestingly, our method also implicates that SoxS interactions at -55, where it serves as a repressor of our promoter, are stabilizing in nature. Furthermore, this method does not suggest that *−*47 does not have stabilizing interactions, only that the stabilization at *−*43, *−*53 and *−*55 is stronger.

Motivated by our detection of stabilization in the regulatory response of SoxS, which is known to act through stabilization, we applied the manifold analysis to the pairwise regulation of each TF covered in the position sweep. In Fig. 5C, we see that for repressing TFs YqhC, MelR, and GntR the pairwise Fold-Change profiles for select pairwise positions deviate from a straight line, implying a dependence on positive stabilization (*β >* 1). For YqhC, regulation at position *−*53 presented a *κ* value of around 7, indicating significant stabilization that is position-specific (a feature shared by both MelR and GntR) and is co-incident with a position of curvature for the SoxS activator. This feature suggests that despite opposing regulatory outcomes, both TFs stabilize RNAP at the same position - pointing to a sensitivity of this regulatory mode to binding location. However, the activator MetR presents no evidence for stabilization despite its role in activating expression at positions *−*41,*−*43, and *−*53, which is confirmed by plotting the manifold response at the position of highest curvature in Fig. 5D. Here the manifold more closely resembles repression by NtrC, with profiles that are straight lines. This raises the possibility that kinetic modulation of recruitment and transcription initiation by MetR is fundamentally distinct from SoxS despite both TFs functioning as activators.

To demonstrate the position landscape of stabilization across TFs, in Fig. 5E we plot the *κ*_max_ values for 21 out of the 24 TFs. The full list of values can be found in Table 1. We excluded 3 TFs (IclR, BaeR, and CsgD) from this analysis because either the response functions were a poor fit the thermodynamic model (Fig. S7) or there was only a single regulatory position (precluding assessment of stabilization using the concentration manifold). We note the predominance of GntR enrichment across promoter-proximal binding locations, suggesting this TF utilizes stabilization in a manner that is insensitive to binding orientation (upstream or downstream) and distance from the promoter. The TFs Nac, AgaR, and PdhR each emerge with potential stabilization in their regulatory function across distinct binding positions with significant *κ*_max_ values. We also find that the stabilization presented at proximal binding positions for repressors, also occurs downstream of the promoter, with the +5 position revealing several TFs that repress at that location. Curiously, while the TF Nac activates at position *−*47, we find no evidence for stabilization at that position, but instead find the highest *κ*_max_ value for the TF at its downstream repressing position. These observations highlight that activation is not synonymous with stabilization of RNAP and reinforces the possibility that several repressors may exert their quantitative regulatory function through the mechanism of a kinetic trap [75], rather than through inhibition of RNAP binding at the promoter.

**Table 1:** *κ*_max_ values for positions of greatest curvature from the manifold analysis. The median and interval comprising 86th percentile of the parameter range is reported.

| TF | Binding Position | $\kappa_{\max}$ |
| --- | --- | --- |
| GntR | -41 | $14.306^{+8.761}_{-5.252}$ |
| Nac | 5.0 | $7.752^{+22.344}_{-5.644}$ |
| YqhC | -53.0 | $6.176^{+5.658}_{-3.599}$ |
| PdhR | -41.0 | $4.175^{+0.851}_{-0.859}$ |
| AgaR | -43.0 | $3.914^{+3.261}_{-1.816}$ |
| MelR | -43.0 | $3.238^{+0.319}_{-0.401}$ |
| UlaR | -49.0 | $2.980^{+0.719}_{-0.602}$ |
| TrpR | 5.0 | $2.713^{+0.719}_{-0.497}$ |
| GcvA | -59.0 | $2.655^{+0.467}_{-0.425}$ |
| CueR | -41.0 | $2.382^{+0.681}_{-0.476}$ |
| SoxS | -55.0 | $2.329^{+0.935}_{-0.715}$ |
| AcrR | -49.0 | $1.944^{+0.134}_{-0.0862}$ |
| MngR | -57.0 | $1.694^{+0.483}_{-0.248}$ |
| GlpR | -41.0 | $1.600^{+0.279}_{-0.202}$ |
| BetI | 5.0 | $1.586^{+0.281}_{-0.196}$ |
| NagC | -49.0 | $1.531^{+0.207}_{-0.197}$ |
| NtrC | -41.0 | $1.489^{+0.065}_{-0.121}$ |
| MetR | -47.0 | $1.469^{+0.558}_{-0.408}$ |
| CusR | -41.0 | $1.408^{+0.538}_{-0.447}$ |
| AraC | -43.0 | $1.404^{+0.203}_{-0.146}$ |
| RutR | 5.0 | $1.044^{+0.465}_{-0.452}$ |

### Promoter dependence of regulatory effect

An important result from the manifold analysis was the suggestion that the molecular mechanisms of TFs with diverse net regulatory outcomes (activation and repression), could feature stabilization (*β >* 1) as a mechanism. In Fig 6A, we list the *κ*_max_ values for select TFs that encompass TF-position circuits with noticeable curvature in the regulatory response as well as those without it. The presence of stabilization (*β >* 1) for a given TF/binding position offers an explicit prediction on how regulation should depend on the strength of the promoter as highlighted in Fig. 1E and Fig. 6B. Specifically, we expect that Fold-change increases on weaker promoters for TFs that operate through stabilization, *β >* 1; that is to say repressors repress less while activators activate more. On the other hand, if a TF operates through destabilization, *β <* 1, we expect Fold-change to decrease for weaker promoters; repressors repress more and activators activate less. As such, measuring the promoter-dependence of a regulatory interaction is, in principle, a powerful method to reveal the mechanisms of regulation by a TF. By tracing the inferred *FC*_max_ across promoter strength, we can determine the magnitude of stabilization in the regulatory response (*β*) and infer the magnitude of acceleration (*α*) as a result.

**Figure 6:**
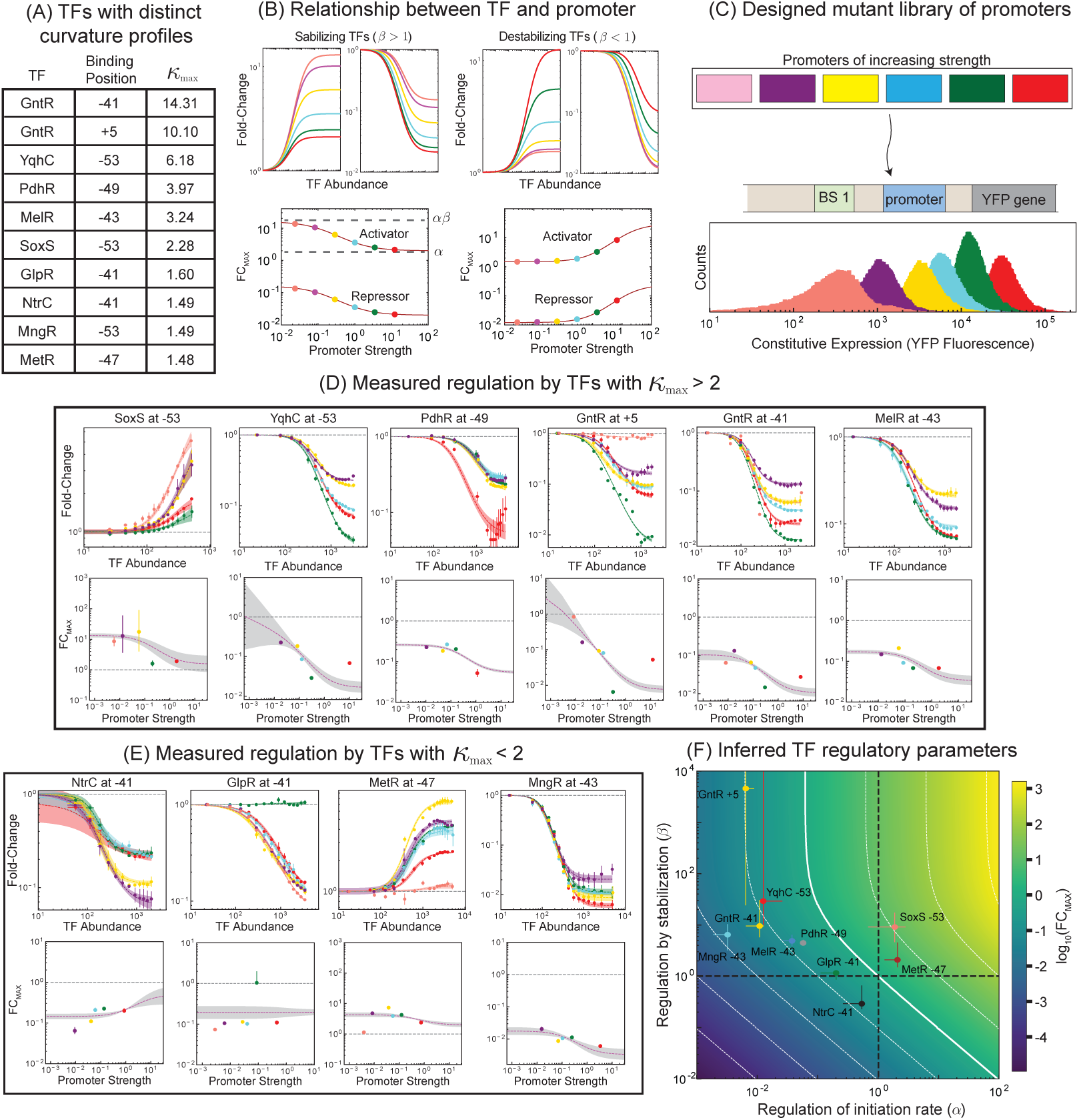
Regulatory function of different TFs on a library of promoters. (A) List of TFs with their respective *κ*_max_ values. (B) We measure regulation by these TFs on a library of *σ*^70^ promoters. (C) *FC*_max_ inferred from the TF abundance input-output function on a suite of different promoters with varying basal occupancy can be used to uncover the role of stabilization and acceleration in the regulatory response. (B) Design of a suite of *σ*^70^ synthetic promoter constructs with different constitutive expression levels. Colors indicate promoter strength. (D) TFs with *κ*_max_ *>* 2 show the expected relationship between *FC*_max_ and promoter strength, *P*. Errorbars in the top row represent the uncertainty associated with fold-change measurement determined by bootstrapping with the shaded area representing the thermodynamic model uncertainty. Errorbars for the scatter plots in the bottom row represent the 64*^th^* percentile interval range of the inferred promoter FC_max_. (E) TFs with *κ*_max_ *<* 2 showed diverse relationships between *FC*_max_ and promoter strength, *P*. (F) The Phase Space of TF regulatory modes as a function of TF identity and binding position. The majority of TFs that regulate exclusively by repression do so with a significant stabilization (*β >* 1) component - with some possessing larger coefficients than activating TFs.

To test the relationship between TFs characterized in the previous section and promoter identity, we designed a set of 6 promoters with constitutive expression levels spanning over a 100*−*fold as shown in Fig. 6C. These promoters were designed by mutating the *−*10 and *−*35 regions of our synthetic promoter through single-nucleotide substitutions (see Methods) to achieve this range in constitutive expressions (Fig. 2). We focused on building the library primarily through mutations in *−*35 as we reasoned these changes would be most likely to alter occupancy of RNAP at the promoter rather than affecting the initiation rate of transcription [76–78].

In Fig. 6D we measure the Fold-change vs TF concentration for distinct TF-position regulatory circuits that demonstrate high *κ*_max_ values from the manifold analysis. Overall we see the same relationship for all of these regulatory interactions: stronger promoters have lower fold-changes. This relationship holds for both the activator SoxS and the four different repressors in this panel. Despite their opposite net function, they have the same relationship between fold-change and promoter strength owing to their shared regulatory mechanism which was predicted from our manifold analysis. Importantly, we do find one case where we measure large curvature in the manifold approach (*i.e.* high *κ*_max_ value), yet we see no evidence for stabilization from the promoter titration response (Fig. S10). Furthermore, in Fig. S9 we test CpxR at *−*64, which we previously had determined to be stabilizing (*β >* 1) and accelerating (*α >* 1) and find that the promoter titration supports these findings [14].

In Fig. 6E we test several regulators where we did not find strong evidence of stabilization from our manifold approach. In these cases, we expect one of several possible outcomes: the TF does stabilize at that position but less than or similar to the other positions we measured in the manifold, the TF does not stabilize (*β* = 1) or actively destabilizes RNAP (*β <* 1). Testing the regulation of this cohort with *κ <* 1 on promoter collection, we find a clear relationship between *FC*_max_ and promoter strength for NtrC regulation at *−*41, with the stronger promoters being less repressed as compared to the weaker promoters for NtrC (Fig. 6E). This is the opposite relationship we saw previously for TFs with *κ*_max_ *>* 2 and is in line with our model predictions for destabilization. For GlpR, we see no relationship between *FC*_max_ and promoter strength, demonstrating repression that does not appear to function through stabilization and instead functions on initiation rate (*α <* 1). MetR at *−*47 is less clear but perhaps demonstrates a weak stabilization profile, with a stronger dependence on modulating transcription initiation to generate the activating response at that position. The regulation of MngR at position *−*53, demonstrates the strongest stabilizer among these TFs with low *κ*_max_ values.

Finally, using this approach we can infer the value of both regulatory parameters (*α* and *β*) for the specific TF-position regulation. We find these values by fitting the theory in Fig. 1B to the plots of *FC*_max_ as a function of promoter strength (dashed lines). In Fig. 6F, We show a parametric plot of regulatory function by plotting *β* and *α* pairs for each characterized regulatory interaction (parameter values listed in Table 3). We find that incoherent regulation (up-regulating one step while down-regulating another) seems to play a major role for regulators examined here, suggesting a potential rule that delineates a relationship between the modes of acceleration and stabilization for these *E.coli* TFs.

**Table 2:**
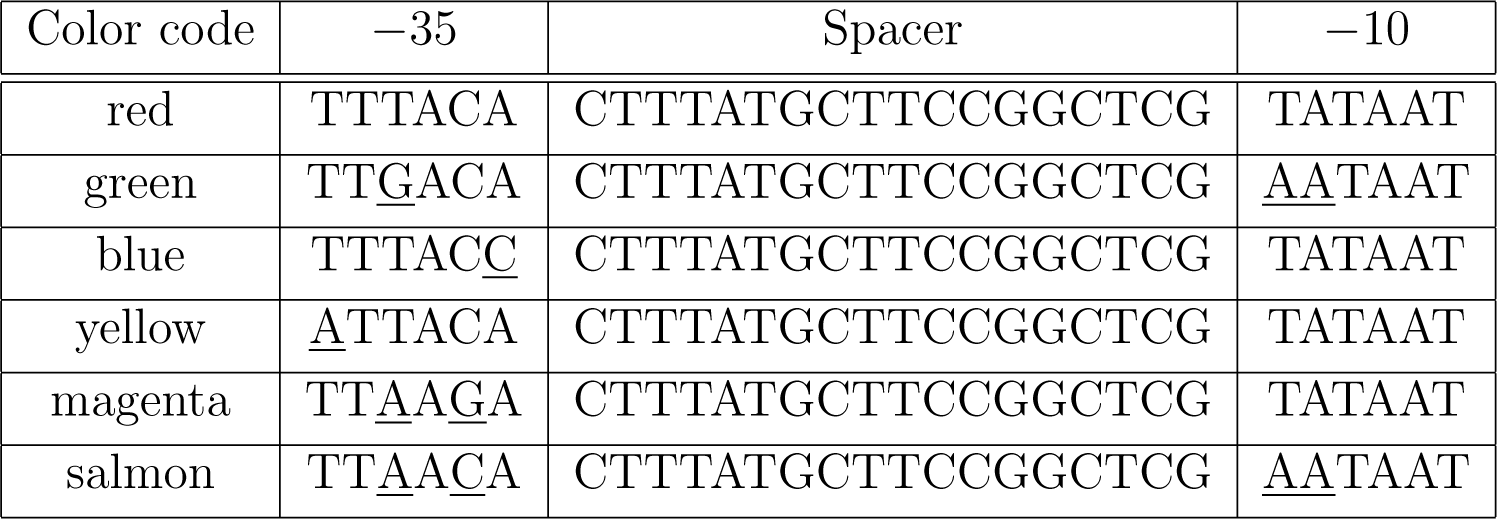
*σ*^70^ promoter sequences used in synthetic circuit construction. Six promoter sequences with mutations in the *−*35 and *−*10 hexamers provided sufficient dynamic range to test the impact of TF regulatory function on different RNAP effective concentration levels.

**Table 3:** Median and. 86*^th^* percentile interval range for *log*_10_*α* and *log*_10_*β* across 10 designed TF-position-sequence circuits.

| TF | Binding Position | $\log_{10}\beta$ | $\log_{10}\alpha$ |
| --- | --- | --- | --- |
| YqhC | −53 | $1.465^{4.840}_{1.004}$ | $-1.900^{-1.585}_{-1.949}$ |
| MelR | −43 | $0.688^{0.811}_{0.588}$ | $-1.423^{-1.362}_{-1.574}$ |
| GntR | −41 | $0.98022^{1.4832}_{0.7606}$ | $-1.962^{-1.889}_{-2.182}$ |
| GntR | +5 | $3.660^{6.037}_{1.383}$ | $-2.190^{-2.047}_{-2.199}$ |
| NtrC | −43 | $-0.5357^{-0.178}_{-0.639}$ | $-0.270^{-0.250}_{-0.580}$ |
| SoxS | −53 | $0.961^{1.240}_{0.6664}$ | $-0.162^{0.4553}_{0.2760}$ |
| MetR | −43 | $0.316^{0.659}_{0.171}$ | $0.317^{0.372}_{0.105}$ |
| GlpR | −43 | $0.065^{0.099}_{-0.061}$ | $-0.6983^{-0.658}_{-0.950}$ |
| MngR | −43 | $0.809^{1.014}_{0.507}$ | $-2.489^{-2.4697}_{-2.733}$ |
| PdhR | −43 | $0.652^{0.711}_{0.643}$ | $-1.238^{-1.218}_{-1.289}$ |

## Discussion

Our ability to interpret promoter sequences is predicated on understanding the sequence-rules inscribed in regulatory DNA. Systematic efforts have been developed to probe the sequences TFs interact with that sets the occupancy landscape of these factors yet less is known about the specific role TFs play in modulating distinct steps of the transcription cycle. This presents a significant knowledge gap, as expression levels of a regulated gene are determined by more than just the presence or absence of a TF binding sequence or motif, but emerge from the interplay of TF binding sequence, location, and the basal occupancy of the promoter in the absence of regulation [39, 40, 79, 80]. These and other features, collectively referred to as promoter context, present a central challenge in generalizing findings of TF regulatory activity across different promoters and genetic backgrounds.

In this study, we examine a handful of genetic determinants using synthetic gene circuits to isolate the relationship between these features and the TFs ability to regulate two steps of the transcription process: recruitment of RNAP to the promoter sequence (stabilization - *β*) and initiation of transcription (acceleration - *α*). We profile 90 *E.coli* TFs, (roughly 1/3 of annotated TFs) and isolate the contributions of TF binding sequences and binding positions to a TFs ability to modulate these steps. Despite the limited number of binding positions and sequences tested per TF, we find examples in our data that demonstrate both TF operator sequence and operator position shape TF regulatory function (*α* and *β*). We also find certain TFs that regulate at distal (*−*103), proximal upstream (*−*57) and downstream (+1) positions with the same degree of regulation, suggesting TF-specific regulatory principles that could be useful in applications to design gene regulatory circuits with tailored regulatory outcomes.

As stabilization of RNAP is known to occur for several well-characterized *E.coli* TFs [33, 81, 82], we sought to infer the degree to which this mode is utilized for a subset of TFs that we profiled. Strikingly, we found evidence for this regulatory mechanism across several TFs including those that function exclusively as repressors in the positions tested. This suggested a conserved mode of TF activity where RNAP-stabilizing interactions appeared to be a driver of fold-change. Stabilization was consistent for certain repressing TFs across upstream and downstream binding position and provides evidence of repression via a kinetic trap (strong recruitment *β >* 1, reduced initiation *α <* 1 [75, 83, 84]), as the dominant mode of negative gene regulation in our data. The degree to which we uncovered this dependence on stabilization, was unexpected, as repression by kinetic competition (*β <* 1) is a well-established paradigm of gene regulation [80], and shapes much of how we view repression as hindering RNAP recruitment to promoters. Here we do not see clear evidence of kinetic competition, except perhaps weakly for NtrC, which manifests as reduced repression on strong promoters compared to weak promoters.

The extensive use of stabilization by TFs of diverse net regulatory outcomes suggests a conserved relationship between TF function and the promotoers they regulate in line with predictions from a thermodynamic model of gene regulation [11, 14, 41, 85]. By quantifying the mechanisms that drive regulation for individual TFs, we can more easily understand the differences in regulation across distinct natural promoters. The approach highlighted here using systematically designed promoters, provides a powerful tool to characterize TFs in terms of their regulatory mechanisms (stabilization and acceleration) while also providing testable hypotheses on their effect in novel promoter contexts. Furthermore, the applications to dissect combinatorial gene regulation are also worth considering; the mechanisms by which TFs function should also dictate their ability to work in tandem [10, 13]; models that consider the mechanisms by which fold-change is enacted rather than simply the net effect itself should be capable of predicting the regulation of promoters with higher complexity. Together, this study uncovers the important connection between the regulatory function of TFs and the *cis*-acting determinants that interpret and transfer this function into quantitative gene expression profiles.

## METHOD DETAILS

### Selection of TF binding sequences from RegulonDB

The TF binding sequences used in interrogating the role of position and sequence on TF regulatory modes were curated from RegulonDB [47]. We performed an initial inventory of TFs in our synthetic library and determined which of these TFs overlapped with detailed information on TF binding sequence in this database and were amenable to our cloning procedure described in this section. TFs BirA, CecR, DicA, DnaA, HigA, LexA, MazE, MqsA, PdeL, YefM, OxyR were excluded from our analysis as their respective sequences returned from our curation (described following) contained the recognition site for the BbsI enzyme used in cloning the synthetic circuits. We settled on 90 TFs with information on binding position of TFs throughout the genome, their qualitative regulatory function (activation or repression), the degree of evidence for binding and regulation, and their nucleotide sequence. We then filtered the sequences to promoters that were bound by *σ*70 and selected the two (or three) binding sequences. For TFs with many possible binding sequences after our initial curation, we constructed a position weight matrix based on the available sequences and ranked each sequence based on similarity to the matrix. The two highest-ranked sequences were selected for cloning into the synthetic gene circuits described in the Results.

### Culture Conditions for Measurements of TF Regulatory Function

We measure the regulation of *E.coli* TFs on synthetic circuits with different TF binding sequences, binding positions, and promoter strengths using an inducible transcription factor library [41]. Specifically, these TFs had their coding region removed from their endogenous genomic loci. and replaced in the *ybcn* locus. The TF coding sequence (fused via a linker sequence to mCherry) at this locus is regulated by the *pTet* promoter allowing for inducible expression. All library strains harbored the TetR cassette integrated into the *gsp1* locus expressed from a constitutive promoter. To measure the effect of a TF on a given synthetic circuit, we grew the strains in LB media overnight, then made a 1:20000 dilution in M9 minimal media. All cultures were grown in a shaking incubator (250 RPM. 37*^◦^*C) in 96-well plates for reproducibility, and we aimed to measure cultures once they reached steadystate (approximately OD600 = 0.1 *−* 0.2). To vary the input TF concentration, we cultured each of the strains with the addition of aTC (anhydrous tetracycline) at the following concentrations - 0ng/ml, 0.75ng/ml, 1.5ng/ml, 2ng/ml, 4ng/ml and 6ng/ml. We found these aTC conditions provided the best dynamic induction of TFs under the *pTet* promoter. After the target OD, was reached, strains were diluted in M9 minimal media, and the mCherry and YFP fluorescence were measured via flow cytometery (BD LSRFortessa X-20- Model #: 656385).

### Cloning of synthetic gene circuits with defined regulatory architecture into respective TF library strains

To clone 180 TF binding sequences at three binding positions in the synthetic circuits, we ordered the TF binding sequences as single-stranded DNA oligos and annealed them to their reverse complements in a predefined arrangement in a 96-well PCR plate. The oligos had overhangs that were designed for digestion by the Type IIs BbsI restriction enzyme with the following sequence ‘5’-GAAGACXXCGTG-3” to facilitate cloning into a set of plasmids developed previously [14]. These plasmids have the *ccdB* cassette integrated at defined positions relative to the TSS with overhangs for the same enzyme. Cloning of the TF binding sequences at a given position was done in a single digestion and ligation reaction by incubation with BbsI and T7 ligase with the addition of isolated position construct plasmids. We performed temperature oscillations between 37° and 16° at 5-minute intervals for optimal digestion and ligation for 30 cycles. The reaction mixture was then transformed into chemically competent cells, cultured in SOC media for 1 hour, and plated on agar plates. PCR sequencing was done to confirm the incorporation of the oligo for all wells using a common priming site flanking the promoter region. Select clones from the 96 well plates were sent for Sanger sequencing to verify inclusion of the TF binding sequence at the defined position. We arrayed cultures from the TF library strain in 96 well plates that matched the arrangement of the cognate TF binding sequences. We then mini-prepped the TF-binding sequence/position plasmids in 96 well plates, isolated the DNA through ethanol/isopropanol precipitation, and transformed into chemically competent TF library strain cultures. The competent cells were generated in 96 well plates, by alternating cycles of washing the cell pellet in CaCl_2_ and centrifugation to remove the supernatant.

### Designing Promoter Sequences to Titrate Basal Promoter Expression

To assess the relative contributions of stabilization and acceleration (*α* and *β*) across several TFs, we designed promoters with varying degrees of basal expression spanning several orders of magnitude. The promoter used in our throughput gene circuits was a variant of the *LacUV5* promoter, the natural promoter that regulates the *lac* operon. Based on prior work testing different variants of this promoter [86], we used 4 variants from this work, with two additional promoters designed from considering the sequence energy matrix of the Sigma70 promoter as investigated in [78]. These mutations encompassed bases in both the *−*35 and *−*10 sequences: we made no changes to the spacer nucleotides as well as the sequences between the *−*41 to *−*35 and the *−*10 to +1 regions. The sequences are listed in Fig. 2.

### Data Quantification and Statistics

#### Flow cytometry measurements of TF binding position, sequence, and promoter gene regulatory circuits

Measurements of synthetic circuit output as a function of TF abundance were done in 96-well flat-bottom plates via flow-cytometry. Events from a given well were initially gated at data acquisition using a threshold procedure that incorporated the Forward Scatter Area and YFP fluorescence. The event data from a given well were then processed using a custom procedure to refine the number of events that were ultimately used in the calculation of the Fold-Change in promoter expression as detailed in [14]. To compute the Fold-Change in gene expression from the measured values, we measured TF abundance (mCherry) and gene expression (YFP) from the regulated circuits and from circuits with no binding site for the TF (growth rate control). All fluorescence values were background subtracted from measured autofluorescence strains (KO strains of the respective TFs). For measurements of the response functions of 90 TFs, we note that 12 of the TFs did not have appreciable growth in the KO strains in our culture conditions (M9 minimal media supplemented with glucose). For these strains, we used the median of the distribution of the autofluorescence across 78 TFs as a representative measurement as the distribution was fairly tight around the mean Fig. S2.

The mCherry values from these strains were pooled, and events (designated by a vector of cytometer channel values) were binned using the mCherry values. The data was binned proportionally - each interval had approximately the same number of events, with the interval length in mCherry different between bins to accommodate this strategy. The median YFP values associated from a given bin was then normalized to the YFP values from the first bin (fold-change) for both the regulated and no-binding-site strains. The resulting Fold-Change values for the regulated strain at each RFP bin were then normalized to its corresponding Fold-Change value for the no-bindingsite strain in the same bin. For position regulation measurements involving the 24 TFs in Fig. 4, the error bars for a given fold-change measurement were computed using error propagation. This calculation incorporated the variance in autofluorescence background, regulated, and unregulated YFP measurements and was done using custom Python code.

#### Model Evidence Computation to assess the relative contributions of TF binding sequence and location to Regulatory Function

To evaluate the model evidence associated with the contributions of TF binding sequence and location to regulatory function (*α* and *β*), we employ Bayes theorem to evaluate the probability of a thermodynamic model that considers all response functions associated with a given feature (*i.e* binding sequence or position) having a shared FC_max_ parameter (independent model) and an alternate model that has a unique FC_max_ parameter for each response function (dependent model). To describe the process another way, we consider a model where only changes to the binding affinity of the TF (found in *χ*) explain the data without the need to invoke a change in its regulatory function. As we measured the regulatory function of the TF at 3 binding locations (+1, *−*57, *−*103) and and at most three binding sequences (*Seq_a_*, *Seq_b_*, *Seq_c_*), the vector of parameters associated with each model is represented by *θ*:

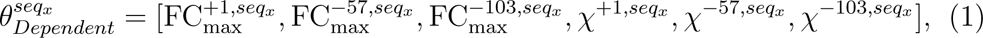

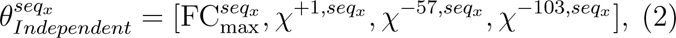

for the hypothesis evaluating position dependence. The subscript x denotes a specific sequence. For sequence dependence, we have the following vector of parameters:

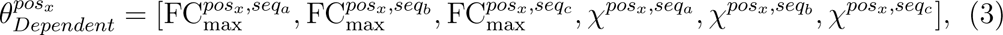

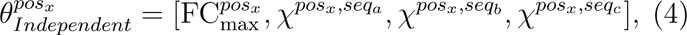

Our goal is to calculate the model evidence *P* (*M |D*) for the dependent and independent models of a given feature (sequence or position). We use Bayes Theorem to delineate this quantity looking at the position feature as an example:

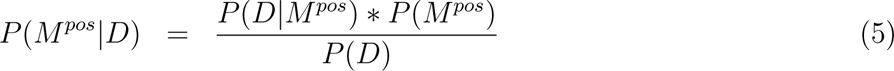

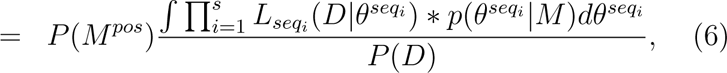

with the variables M and D representing the space of possible models and data. Here, the subscript i represents a given TF binding sequence with s being the total number of sequences profiled for a particular TF. The function *p*(*θ^seq^^i^ |M*) represents the prior distribution of the parameters and *L_seq_* (*D|θ^seq^^i^*) is the likelihood of the data modeled as the product of individual Gaussians:

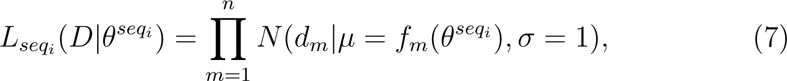

where *d_m_*is an individual data-point and *f_m_*(*θ^seq^^i^*) is the model evaluation that corresponds to that point. To compare the degree to which we favor the independent or dependent model, we compute the Bayes factor which is the ratio of the model evidence

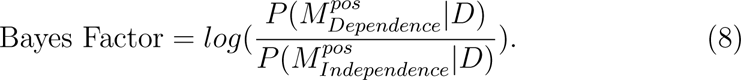

Assessing the model evidence, therefore, requires evaluating an integral over the parameter space which becomes prohibitive when the number of parameters is non-trivial. This makes calculating the posterior model distribution *P* (*M |D*) (model evidence) intractable. To circumvent this problem, we adopt an Approximate Bayesian Computation approach to estimate the posterior model distribution *P* (*M |D*)using sample draws from *p*(*θ^seq^^i^ |M*). Candidate draws are evaluated by approximating *L_seq_* (*D|θ^seq^^i^*) with a dirac delta function:

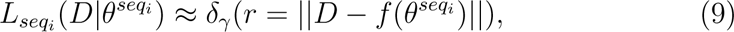

where *r* is the sum of squared residuals of the model and *γ* is a cutoff value that is determined empirically by sampling the distribution of residuals. More information on how *γ* was selected can be found in **Cutoff selection for Approximate Bayesian Computation**. This ensures that only sampled parameters that generate model predictions close to the data are retained and used in computation of the model evidence. We settled on an effective sampling approach that is detailed in the section **Initializing the Prior Distribution**. Once parameters are drawn from the prior distribution, the residuals are computed by evaluating the proposed model to the data. The value of *L*(*D|θ*) is set to 1 if the residuals fall below the threshold and 0 if the residuals are greater than the threshold. Estimation of *P* (*M |D*)) is then reduced to a weighted sum of the prior distributions associated with the parameters of the model.

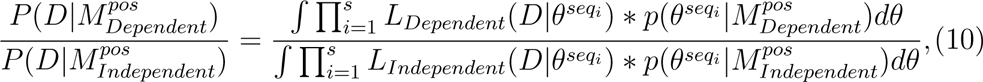

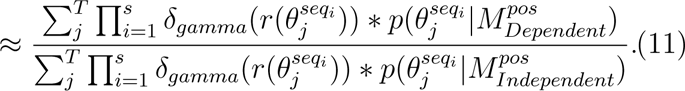

This is where model discrimination manifests: the dependent model - having an independent FC_max_ parameter for each curve, can theoretically attain the smallest residuals from sampling. However, the inclusion of the parameter necessitates an additional weight found in 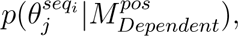 making the increments smaller relative to the independent model. A Bayes Factor larger than 1 would indicate the dependent model outperformed the independent one and implies that the promoter feature under consideration (sequence or position) is essential in setting the regulatory modes (*α* and *β*) of the TF. We ran the model evidence simulation for 100000 runs for each feature for a given TF and report the results in Fig. 3.

### Cutoff Selection for Approximate Bayesian Computation

Estimating the model evidence for the role of TF binding sequence and position in setting stabilization (*β*) and acceleration (*α*) was done using Approximate Bayesian Computation (ABC). This approach provides an approximation to the likelihood function *P* (*D|M*), by simulating parameter draws from the prior distribution *P* (*θ|M*) and determining if the resulting model is a reasonable approximation to the data (see Section **Model Evidence Computation to assess the relative contributions of TF binding sequence and location to Regulatory Function**). We use a cutoff value for the residuals that is determined by considering the distribution of residuals for the independent model:

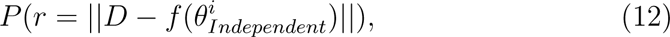

where *s* represents the feature subset (TF binding sequence or position). We consider the independent model distribution as the reference due to the fewer number of parameters in the model that places an upper bound on the minimum of the residual landscape. We can view this in the context of our measurements for the 90 TFs, where we profile 2 binding sequences (designated *Seq_a_, Seq_b_*) at 3 binding positions (+1 *−*57, *−*103). *Seq_a_* has 3 response functions (binding positions) associated with it, and a single draw from the prior *P* (*θ^seq^^a^ |M_independent_*) would provide the parameters for the independent model *f* (*θ^i^*_*Independent*_) to evaluate the residual. To evaluate the role of binding position on FC_max_ (conditioned on *Seq_a_*) we sample 50000 draws from 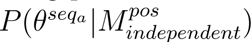 to construct the empirical residual distribution for the given TF feature (see Fig. S4). We find the residual value that corresponds to 2% of the AUC, and used this value as the cutoff (*γ_a_*). The same procedure is done for *Seq_b_* to assign a separate cutoff (provided the associated response functions meet the threshold for regulation), *γ_b_*, allowing us to evaluate the corresponding likelihood functions and perform ABC.

### Initializing the Prior Distribution

Selection of the prior distribution is a crucial step in the approximate Bayesian Computation step. An inappropriately specified distribution function might hamper effective sampling, leading to problems in the distribution of residuals for the independent and dependent models. This would propagate into the calculation of the model evidence leading to a false estimation of the importance of TF binding sequence and position to regulatory function. One possible way to model the prior is as a uniform distribution (assuming *a priori* ignorance on the structure of the parameter set) over a bounded sample space. However, this results in inefficient sampling, particularly when the number of parameters grow (the curse of dimensionality). In the context of our hypothesis, the thermodynamic model associated with a given *cis*-regulatory feature (TF binding position or sequence) has a higher dimensional space for the feature-dependent model.

To address this, we construct the prior distribution by running an inference algorithm over the joint distribution of the parameter space (*p*(*θ|M, D*)) and initializing a kernel density estimate using the markov chains from the parameter runs. This places an effective constraint on the parameter space, allowing for a distribution of residuals that captures the added parameters in the Dependent model while preserving the joint structure of the sample space across the associated parameters (see Fig. S3, Fig. S4). We built individual kernels for each of the 28 TFs considered and drew samples from these kernels during their respective model computations. MCMC sampling was carried out using the PyMC3 probabilistic module in Python. We initialized 4 separate chains, with parameters runs of 5500 draws per chain for each feature.

#### Assessing periodicity in the regulatory response of upstream TF activity

The 9 upstream response functions for the 24 TFs measured from *−*41 to *−*59 bps relative to the TSS were analyzed to determine the dominant frequencies associated with the regulatory response (Fig. 4C). To account for positions of no regulation, we used the Lomb-Scargle algorithm to generate a periodogram from data sampled on an uneven interval [87]. We find a clear peak in the angular frequency space corresponding to 10.94bps (one helical turn of DNA) when looking across the regulatory function of the 24 TFs, with a second smaller peak at around 6bps.

#### Derivation and Fitting of the Manifold Model

To uncover the role of stabilization in the regulatory response of 24 TFs, we extended the manifold model approach detailed in [14]. Briefly, this approach considers the relationship between the Fold-Change at two distinct binding positions, using thermodynamic formalism. By considering the TF binding affinity equal between the regulatory positions, we arrive at an explicit relationship between the fold-changes that depends purely on the regulatory function (*α* and *β*) of the TF at these positions. While the results of Fig. 3 suggest that several TFs evaluated at the 3 binding positions are consistent with a model in which the binding affinity is altered to account for changes in the TF response function, we note that all but one of the 3 binding positions (*−*57) are represented in the position sweep. This allowed us to consider the manifold approach for all of the 24 TFs. In contrast to the previous approach. we relaxed a critical assumption from the previous iteration that assumed the reference position operated through destabilization (*β*). This allowed for the possibility of no or appreciable stabilization at the reference position. As stated in the previous reference and main text, the manifold reformulation of the thermodynamic model yields 4 free parameters: the maximal fold-change corresponding to the two regulatory positions under consideration: 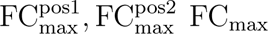 and the curvature associated with the response (*κ*). Our primary objective using this manifold approach is to determine the value of *κ* for a given pairwise position comparison, as this allowed for an assessment of the possibility that a regulatory factor at a given bidding position engages in stabilization. However, as we now relax the assumption that the reference position operates through destabilization, it precludes direct inference of the value of the *β* parameter and allows us to ascertain the magnitude of *κ* (ratio of 1 + *Pβ* at one position to another).

To find the value of *κ*, we first fit the position specific FC_max_ for all regulatory positions (see Fig. S7) according to the thermodynamic model. The model was similar to the one used in evaluating the contribution of TF binding sequence and position to gene expression (Fig. 3) as outlined in Fig. 1B. However, we amended the model to include a hill-coefficient as the sharpness of the response across most of the TF response curves required a coefficient *>* 1 (Fig. S8). We then iterate through all pairwise combinations of regulatory positions and binned the pairwise fold-change data according to the manifold procedure. We employ a bootstrapping procedure to estimate the credible interval for the *κ* value associated with the positions under consideration, using the fold-change data across 3 replicates. For each iteration of the bootstrapping, we used a least squares fitting procedure to find the optimal *κ* from the sampled data and report the median and credible interval for these parameters in Fig. S5. In total, we fit N-1 *κ* values using one of the positions as a reference from the regulatory data across *N* positions for a given TF. Each position-specific *κ* value (relative to the pre-selected reference position) was used as a shared parameter for all pairwise manifolds including that position, leveraging the combinatorial structure of the position regulation dataset.

#### Inferring Stabilization and Acceleration Coefficients from Pro**moter Response Functions**

To delineate the role of *α* and *β* in the regulatory response of several *E.coli* TFs at a specific sequence and position, we measured the response functions of designed synthetic circuits where the core promoter strength was tuned by mutations in the -35 and -10 hexamers. The general thermodynamic model outlined in Fig. 1B denotes how the core promoter strength (*P*) influences the fold change profile. As seen in Fig. 6C, the role of *β* is clear - stabilizing TFs (*β >* 1) will have larger fold-changes for the weaker promoters while destabilizing TFs will act in reverse. The role of acceleration (*α*) in the model is evident when we consider the dependence of the FC_max_ parameter as a function of *P*: in the limit of when 1 + *Pβ ≪* 1, FC_max_ is the product of *α* and *β*, while converging to *α* in the limit of 1 + *Pβ ≫* 1. This is intuitive, as when *P* is large, the TFs role in stabilizing RNAP is diminished, and only its role on processes downstream of RNAP recruitment, *i.e* its role in the initiation of transcription - *α*, sets FC_max_.

We first fit FC_max_ and *χ* to the Fold-Change vs TF concentration curves across different promoters by bootstrapping the fold-change from 2 replicate runs. At each resampling of the regulatory profile, an estimate FC_max_ and *χ* is determined using least squares, and the chain of values of across 1000 runs was used to estimate the median for these parameters. We then used the median FC_max_ parameter for each promoter in the FC_max_ vs *P* plots to infer the value of *α* and *β* parameters for a given TF-position-sequence promoter circuit. *P* (effective RNAP concentration) is an inferred thermodynamic quantity in our work and is related to the constitutive expression (*s*) by the following

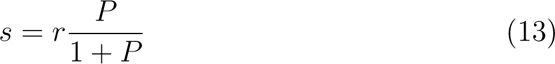

where *r* is the transcription rate. To remove the transcription rate *r*, from consideration, we leveraged the fact that the mutations we made to the *σ*70 hexamers should only affect *P*, and not the downstream kinetic processes of transcription associated with *r*. We therefore used one of the promoters as a reference, and took the ratio of the signal from all other promoters with respect to this. The resulting value of *P* is then given by the relation:

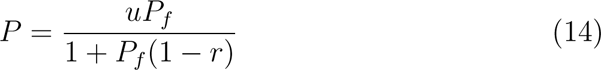

where *u* is the measured ratio of the constitutive expression between promoters and *P_f_* is the reference promoter treated as a fit parameter. As the amount of promoter measurements we made ranged from 4 to 6 promoters, we designed an inference procedure to robustly estimate the interval range for *α* and *β* from this data. We used a bootstrapping approach that would remove either one or two promoter measurements at each iteration, and fit the optimal parameters on the reduced dataset. This approach allowed us to estimate correctly the model expectation at the strong and weak *P* limits, with the median along with 86*^th^* percentile confidence interval for the parameters reported in Fig. 3.

## Supplementary Information

**Fig. S1:**
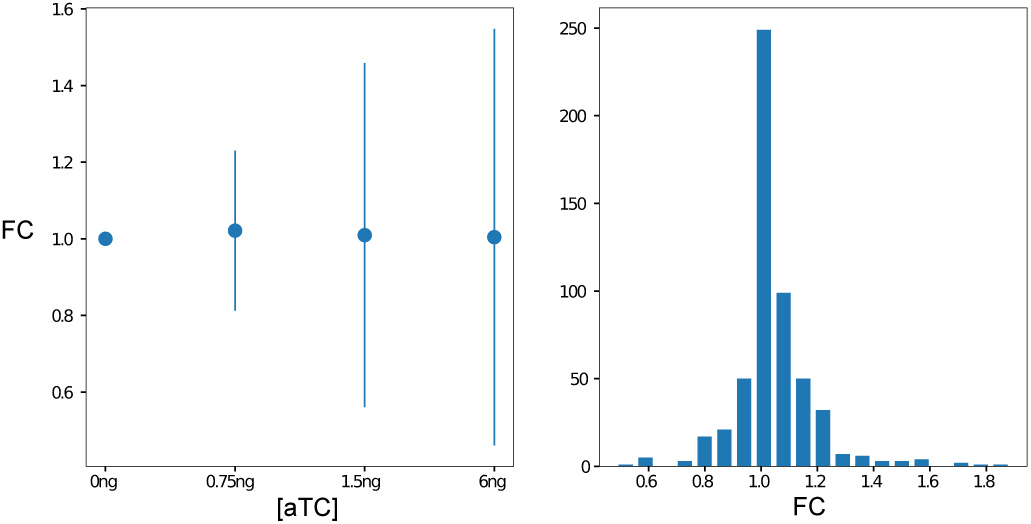
Physiological effects on synthetic circuit readout. The readout from our synthetic circuit (YFP expression) is a convolution of the regulatory architecture (TF binding sequence, location, and promoter strength) as well as global changes in physiology from changing the concentration of the TF. To isolate this effect of cell growth on gene expression, we measured the effect of several aTC induction conditions on the promoter in the absence of any known TF binding sequence with a variant of the *LacUV5* promoter sequence [86]. (A) Normalizing the background subtracted signals to the 0ng aTC concentration, we find the degree to which expression is altered from the unregulated promoter for different concentration of TFs. Error bars represent *±*2*σ*. (B) Histogram combining the fold-changes of the unregulated circuit across 90 TFs at 4 aTC conditions. The large number of physiological measurements gives us confidence in determining synthetic circuits that regulate in Fig. 2

**Fig. S2:**
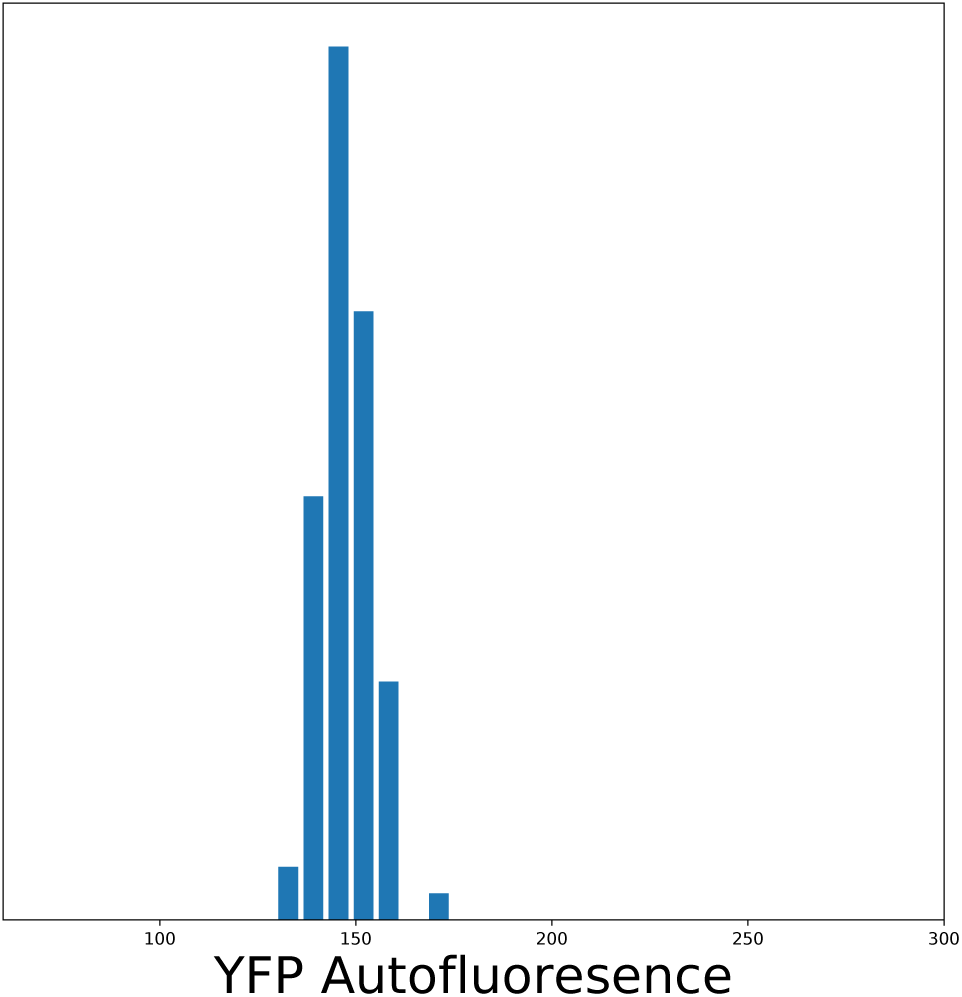
Distribution of autofluorescence (background) measurements across different TFs. To compute the Fold-Change in gene expression described in Methods, we subtracted the measured background fluorescence for each TF library strain using the corresponding KO (knockout) strain. For select TFs (12 of the 90 TFs), the KO strain failed to grow in M9 minimal media. We used the median background fluorescence measured across 78 TFs as the value. As seen, the distribution of autofluorescence values is tightly distributed around the mean.

**Fig. S3:**
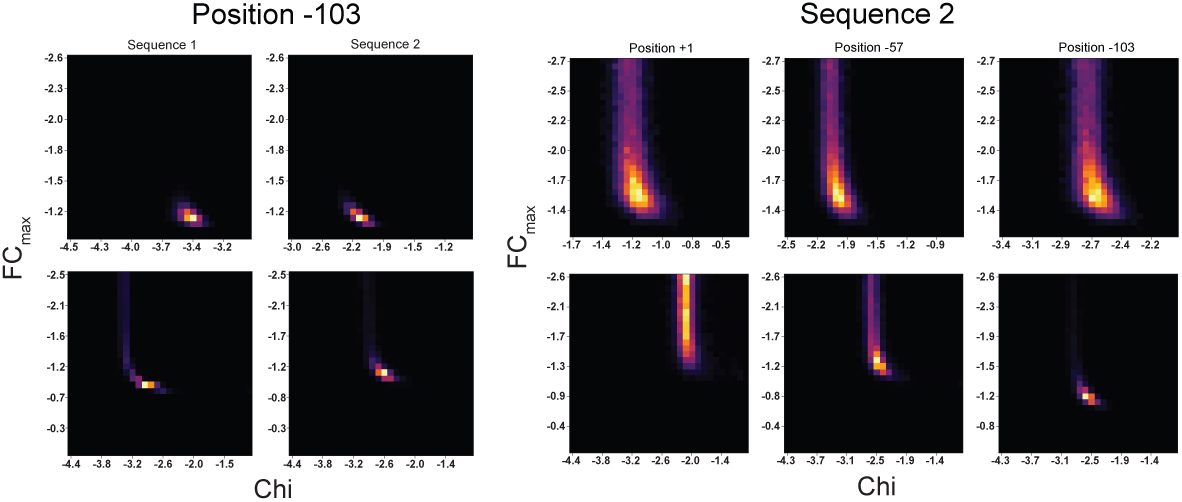
Representative Probability Kernels used in evaluating Model Evidence. To ensure effective sampling of the parameter space in the model evidence computation, we modeled the distribution of prior parameters as a probability distribution derived from inferring the posterior distribution of the two parameters outlined in the general thermodynamic model (FC_max_ and *χ*). The probability kernels for the TF MngR are shown. As only position -103 had both sequences meeting our regulation cutoff, the model evidence for the sequence hypothesis was evaluated only for this position. The top panel is the joint distribution for the constrained model, and the bottom is the free model. Note the change in *χ* in the sampling for the constrained model - signifying the hypothesis that sequence only mediates the binding affinity and not the regulatory function (*α* and *β*) of the TF. For the position hypothesis, Sequence 2 met the criteria for our regulatory cutoff. The constrained model in this instance evaluates the hypothesis that the difference between the regulatory profiles across positions is mediated only through a change in binding affinity of the TF.

**Fig. S4:**
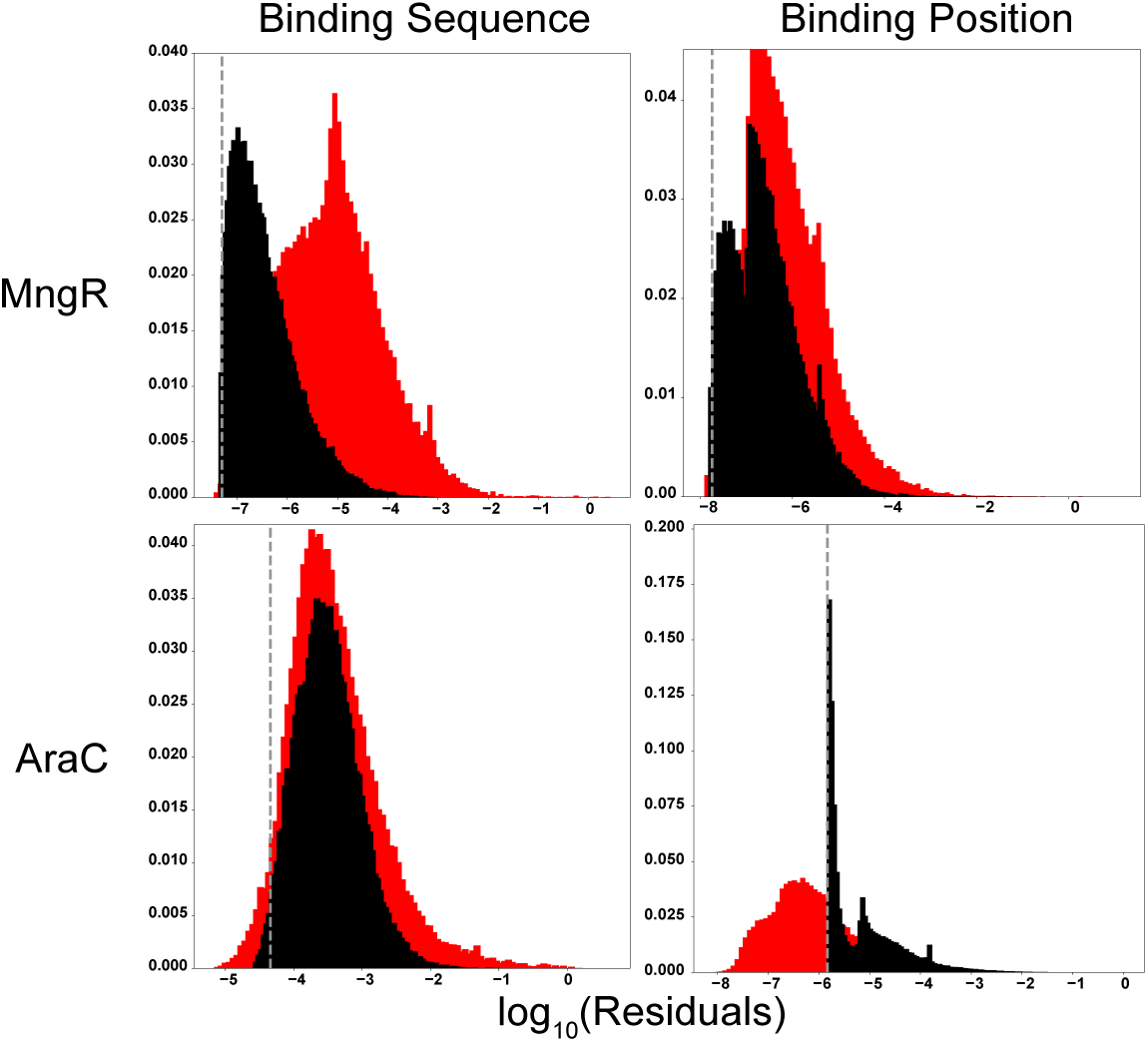
Cutoff selection for Approximate Bayesian Computation of Model Evidence. Plot of the distribution of residuals for the Dependent (red histogram) and Independent (black histogram) models for TFs AraC and MngR. Columns denote the two hypothesis evaluated using the model evidence approach - the role of the tested binding sequences and binding positions in setting FC_max_. Evaluation of the posterior distribution of the model given the data - *P* (*M |D*) - is done using an approximate Bayesian Computation Approach (See Methods Section). To determine if candidate parameter draws from the models are accepted, we considered the empirical distribution of residuals from the constrained model and used a 2 percent threshold. Increments to the posterior model evidence were made if parameter draws for the respective models generated residuals below this threshold. As shown, the sequence hypothesis for both TFs has residuals from both models that overlap significantly. The position hypothesis, however, clearly favors the dependent model for AraC, where there is a discernable shift in the distribution of residuals between the models.

**Fig. S5:**
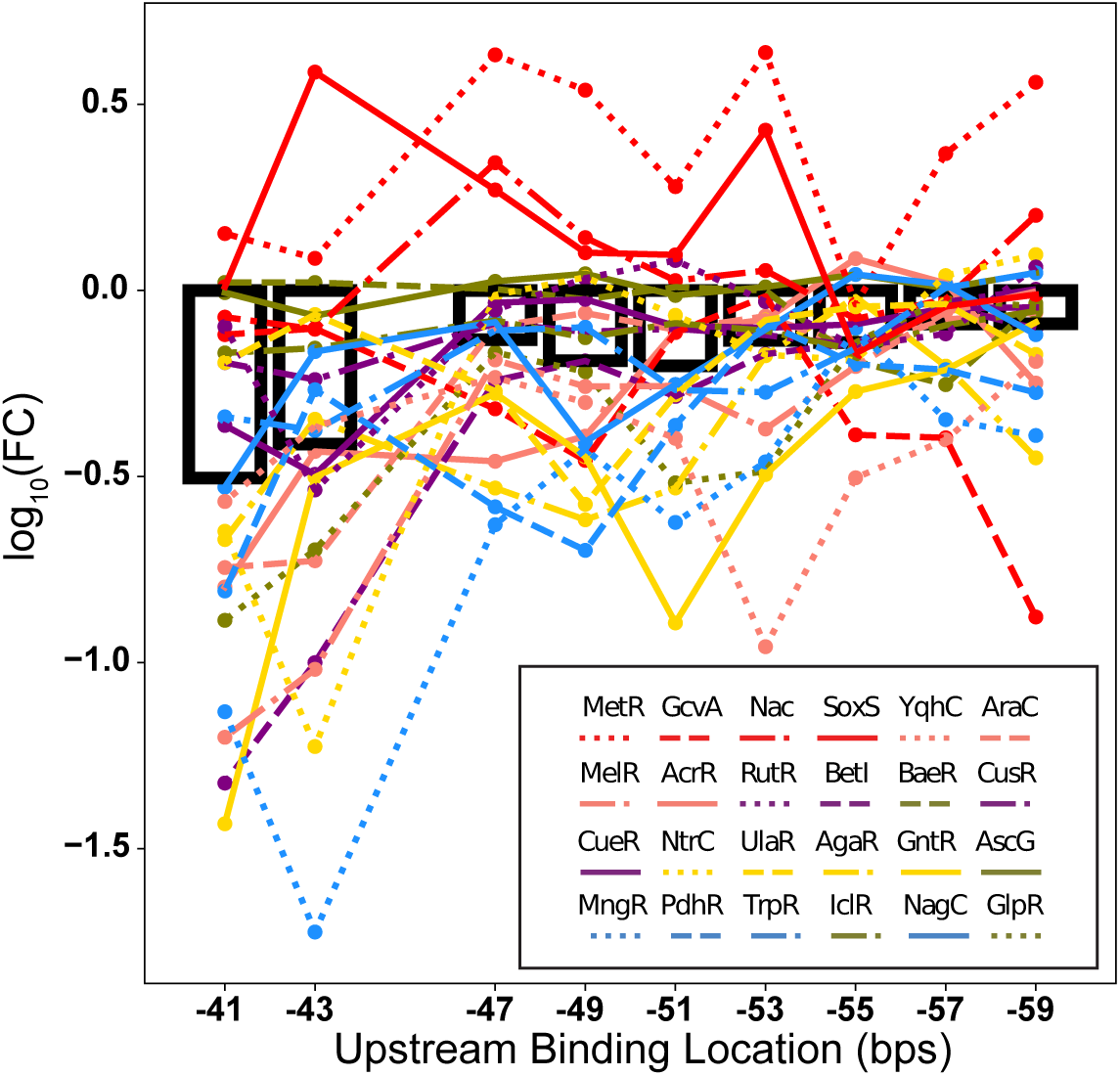
Terminal Fold-Change values for the 9 upstream response functions of 24 TFs measured from. *−*41 **to** *−*59 **bps.** Transcription Factors that activate or repress gene expression tend to adopt a single regulatory outcome over the positions tested with the exceptions for TFs Nac and SoxS. Black bars represent the average regulation at a given position across all the TFs tested.

**Fig. S6:**
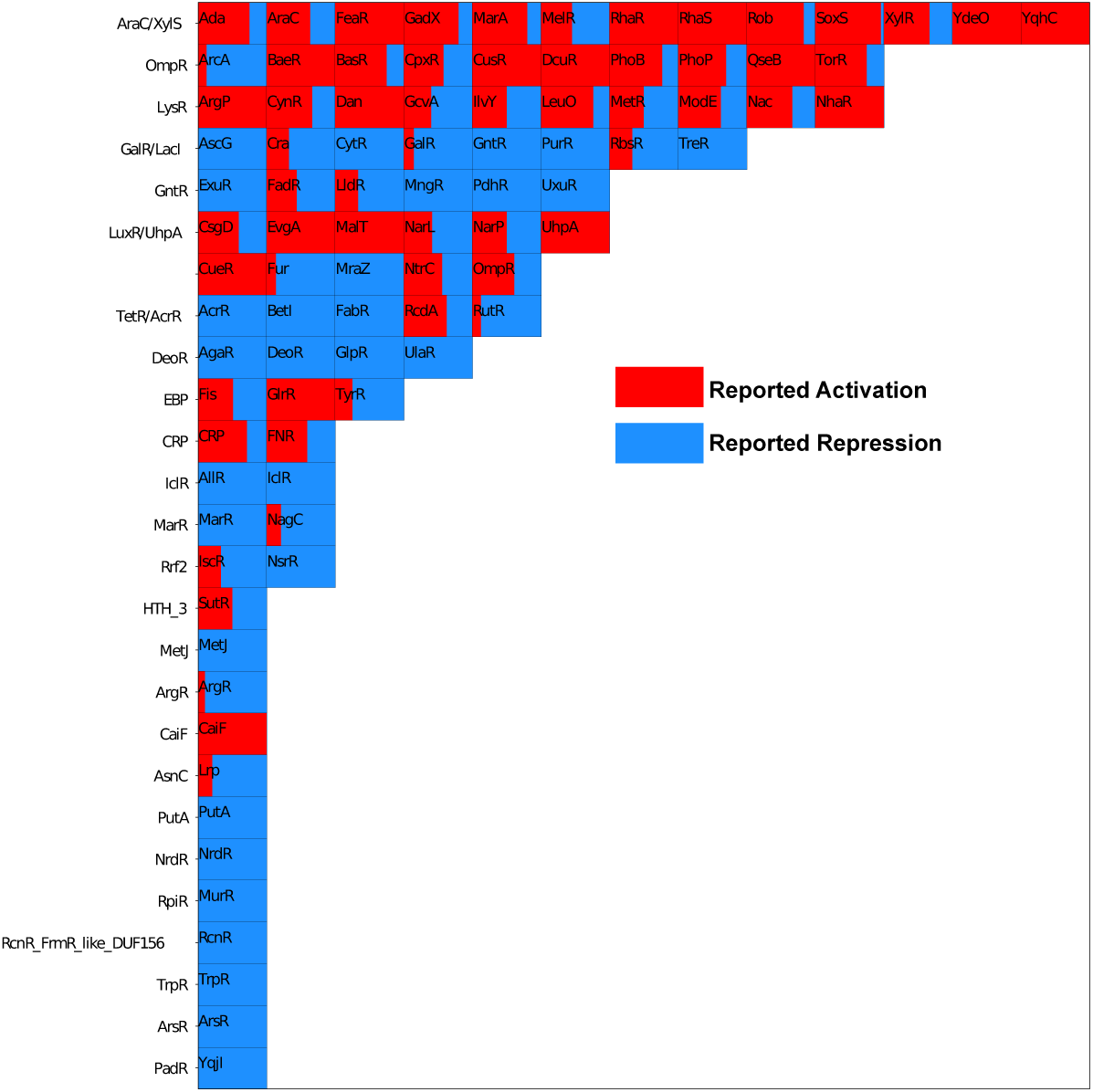
Annotated net regulatory effect on endogenous promoters for the 90 TFs featured in this study. Each square in the plot represents the total number of regulatory interactions ascribed to the TF as compiled from RegulonDB. Shaded areas represent a proportion of these interactions ascribed to activation or repression. Certain TFs such as MetR and GcvA are reported to have dual net outcomes (can activate and repress across different endogenous promoters) while only performing one effect across the regulatory positions tested in our study. We note that the TF YqhC, has only reported activation functions, but we consistently measure repression across multiple positions.

**Fig. S7:**
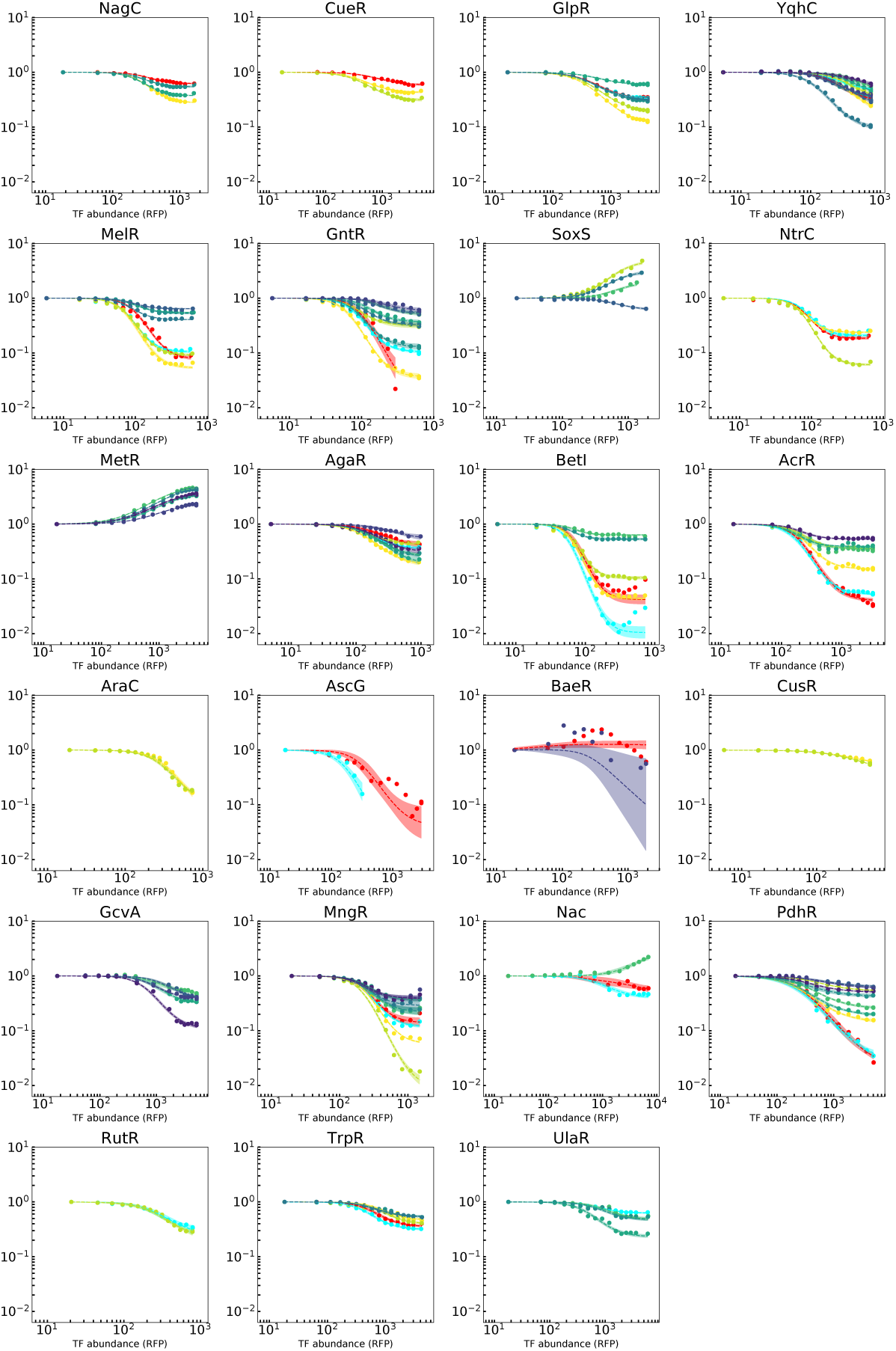
Good agreement between the position-specific measurements of TF regulatory function and thermodynamic model of gene regulation. We fit the measured response functions for the 24 TFs used in quantifying the spatial landscape of regulatory activity to the thermodynamic formalism. Positions with discernible regulation for a given TF are shown. All TFs, with the exception of BaeR, can be well described by the particular thermodynamic model (See Methods Section) used to fit the measured regulatory curves. Note that the TF IclR in Fig. 4E was excluded from the model fitting. Color scheme is identical to the position legend in Fig. 4B. The shaded regions represent one standard deviation from the expected model derived from the bootstrapping procedure detailed in the Methods

**Fig. S8:**
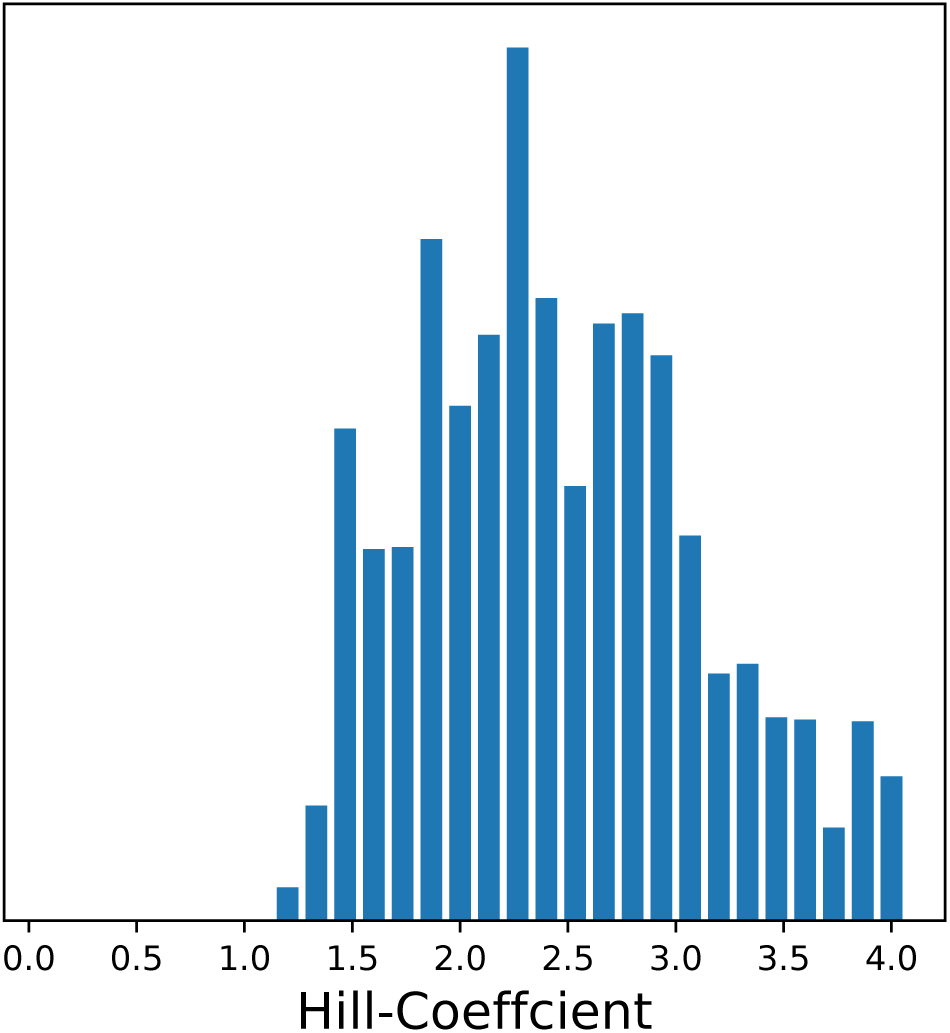
Distribution of Hill Coefficients across the 24 TF position specific regulatory functions. As stated in the Methods, the general thermodynamic model outlined in Fig. 1B was amended to incorporate a hill coefficient to capture the sharpness of each TF response function. To minimize the number of free parameters in the model, the hill coefficient was modeled as a shared parameter across all the position specific regulatory curves for a given TF. The median hill coefficient was 2.35

**Fig. S9:**
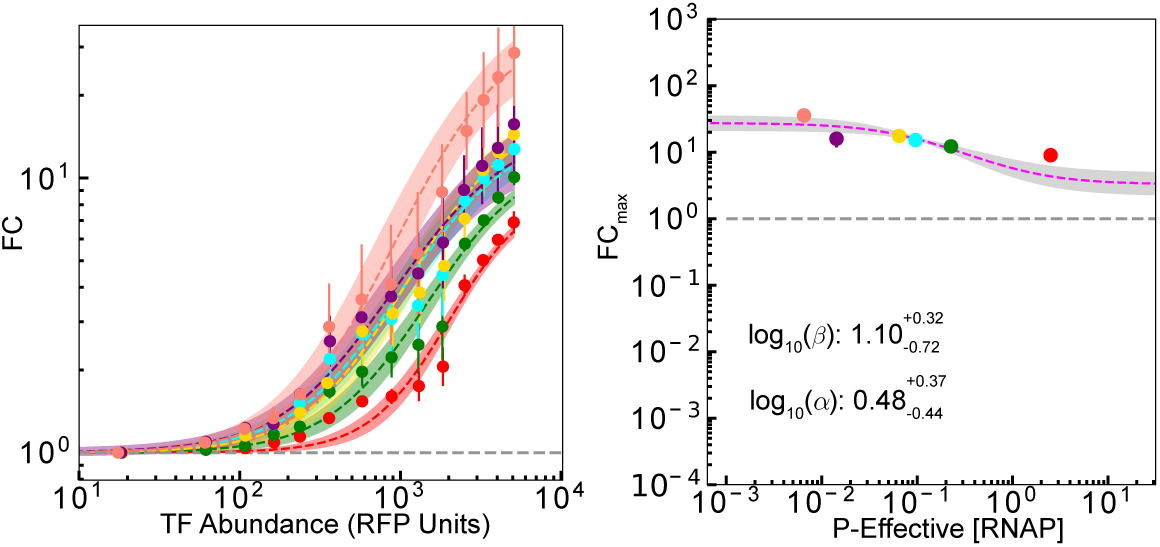
CpxR regulatory activity on promoters of different basal expression strength confirms stabilization in the activation response. CpxR regulation at binding location *−*64 was inferred to engage in stabilization (*β >* 1) using the manifold approach ([14]). To confirm our assessment of this specific mode, we measured the regulatory response of CpxR at this position using synthetic circuits of different constitutive expression levels. We find the characteristic expression pattern of strong activation at weak promoters, with a reduction in activation response at stronger promoters - in line with our expectation of a stabilizing response.

**Fig. S10:**
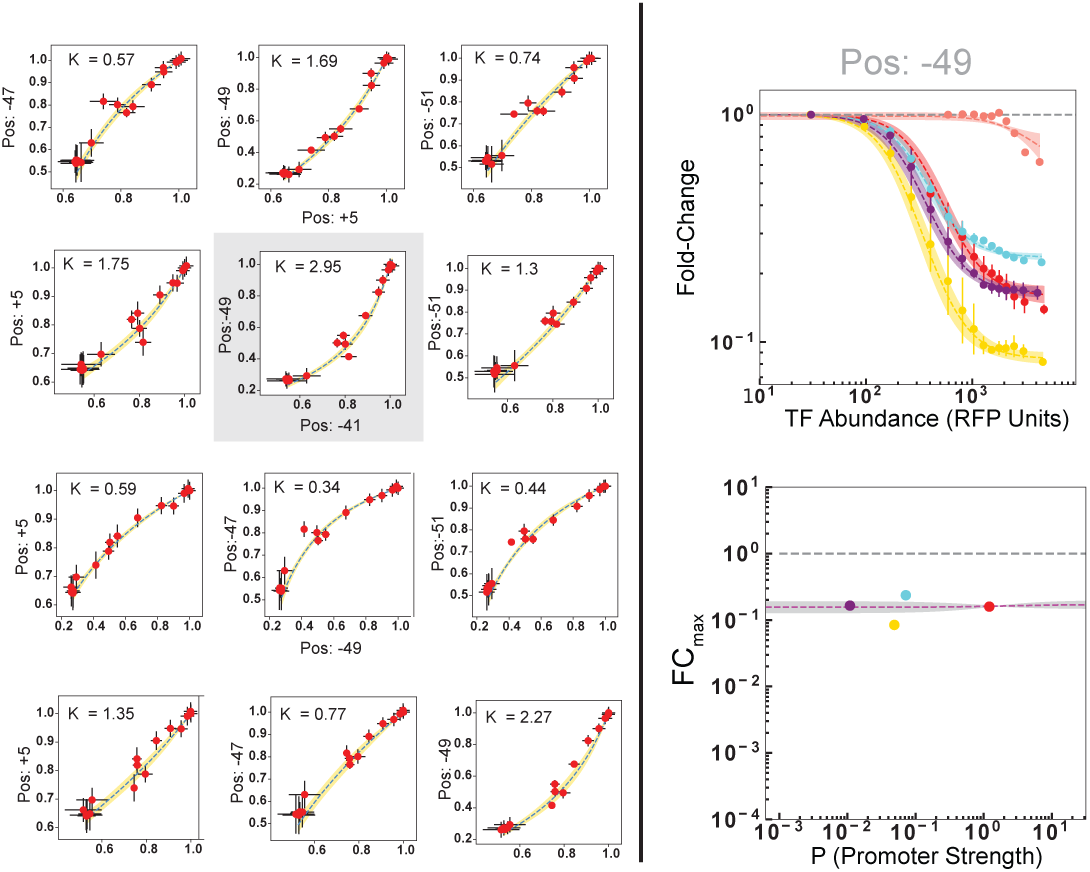
UlaR manifold analysis demonstrates strong curvature - with no evidence for stabilization. UlaR regulation at -49 was inferred to possess a *κ*_max_ = 3.54 - with high possibility for stabilization according to the assumptions laid out in Fig. 5B. However, the regulatory response to circuits of different promoter strengths reveals no underlying evidence for stabilization when plotting the inferred FC_max_ as a function of effective RNAP concentration (*P*). Note the FC_max_ data is across 4 promoters, as the weakest promoter had credible intervals over orders of magnitude and the second strongest promoter (the green data point in Fig. 6C) was not cloned.

